# Multiomics single-cell analysis of human pancreatic islets reveals novel cellular states in health and type 1 diabetes

**DOI:** 10.1101/2021.01.28.428598

**Authors:** Maria Fasolino, Gregory W. Schwartz, Maria L. Golson, Yue J. Wang, Ashleigh Morgan, Chengyang Liu, Jonathan Schug, Jinping Liu, Minghui Wu, Daniel Traum, Ayano Kondo, Catherine L. May, Naomi Goldman, Wenliang Wang, the HPAP Consortium, Michael Feldman, Jason H. Moore, Alberto S. Japp, Michael R. Betts, Robert B. Faryabi, Ali Naji, Klaus H. Kaestner, Golnaz Vahedi

**Author notes:** co-first authors (Maria Fasolino, Gregory W. Schwartz).

## Abstract

Type 1 diabetes (T1D) is an autoimmune disease of only partially defined etiology in which immune cells destroy insulin-producing beta cells. Using single-cell transcriptomics and an advanced analytical strategy to assess pancreatic islets of T1D, autoantibody-positive, and non-diabetic organ donors, we identified both canonical cell types and rare insulin-expressing cells with a hybrid mixture of endocrine and exocrine gene signatures within all donors. We further found elevated expression of MHC Class II pathway genes in exocrine ductal cells of T1D donors, which we confirmed through CyTOF, *in situ* imaging mass cytometry, and immunofluorescence analysis. Taken together, our multimodal analyses identify novel cell types and processes that may contribute to T1D immunopathogenesis and provide new cellular and molecular insights into human pancreas function.

## Introduction

Type 1 diabetes (T1D) is an autoimmune disease which occurs as a consequence of the organ-specific destruction of the insulin-producing beta cells in the islets of Langerhans within the pancreas. This complex disease is characterized by atypical beta-immune interactions including production of beta cell autoantibodies and the immunological attack on beta cells by cytotoxic CD8^+^ T cells (*1, 2*). The immune destruction of beta cells causes deficits in peripheral tissue glucose uptake, increased hepatic glucose production, and hyperglycemia, the combination of which results in devastating effects on metabolic health and severe complications, including blindness, heart disease, and kidney disease (*3–5*).

T1D autoimmunity has been linked to poorly understood genetic and environmental factors. Genome-wide association studies have implicated multiple loci in T1D, with the major histocompatibility complex (MHC) Class II genes as the dominant susceptibility determinant of this disease (*6*). However, the precise cellular context through which T1D susceptibility genes cause the pathogenic destruction of beta cells remains to be discovered. Addressing this question is particularly challenging since the pancreas is a heterogeneous organ, composed of multiple distinct cell types. The exocrine compartment is crucial for nutrient uptake via releasing digestive enzymes, which are produced by acinar cells, into the gastrointestinal tract via the ductal system. Although the endocrine compartment, i.e. the islets of Langerhans, has been the focus of numerous studies using animal models or individual specimens obtained from autopsy collections of human pancreas, the pathogenic relevance of other pancreatic cell types, in particular the exocrine compartment, remains unclear.

Two nontrivial constraints hamper insights into comprehensive identification of the pathogenic cell types in T1D and a better understanding of the initial molecular perturbations that occur during disease pathogenesis: (1) the inability to safely biopsy the human pancreas of living donors and (2) the significant disease progression and beta cell destruction by the time patients are clinically diagnosed with T1D. Therefore, the majority of T1D studies has been performed on peripheral blood leukocytes, which is not the site of pathogenesis. Recently, the Network for Pancreatic Organ Donors (nPOD) (*7*) and the Human Pancreas Analysis Program (HPAP) (*8*) collect pancreatic tissues from deceased organ donors diagnosed with T1D. Additionally, given that many T1D patients harbor beta cell autoantibodies (AAbs) in their bloodstream prior to clinical diagnosis, nPOD and HPAP also collect samples from donors with AAbs towards islet proteins but without a medical history of T1D, in hope of elucidating early pathogenic events. Here, we report an unprecedented multimodal analysis of millions of cells using three high-throughput single-cell techniques in pancreatic tissues of human organ donors collected by HPAP, providing novel cellular and molecular insights into T1D pathogenesis.

## Results

### Single-cell RNA-seq unravels canonical and novel cellular states in the human pancreas

To unmask the molecular perturbations occurring in pancreatic tissues during T1D, we constructed 81,313 single-cell RNA-seq (scRNA-seq) libraries from pancreatic islet cultures of 24 human organ donors representing three categories: individuals with T1D (n = 5), those with AAbs toward pancreatic islet proteins but no clinical diagnosis of T1D (‘AAB+’; n = 8), and those with neither AAbs nor a history of T1D (‘Control’; n = 11) (Fig. 1A and fig. S1 a and b, and table S1). After stringent quality control, we filtered outlier cells, removed doublets, and employed the cell type classifier ‘Garnett’ (*9*) (Fig. 1B, fig. S1, c to e, fig. S2, a to c, and fig. S3, a to e), to cluster 69,645 high-quality cells using ‘TooManyCells’ (*10*), which employs tree visualization to preserve the relationships among cell clusters (Fig. 1C). The resultant classification was confirmed by canonical gene marker expression for each cell type (Fig. 1C and fig. S4, a, c, and d).

**Figure 1:**
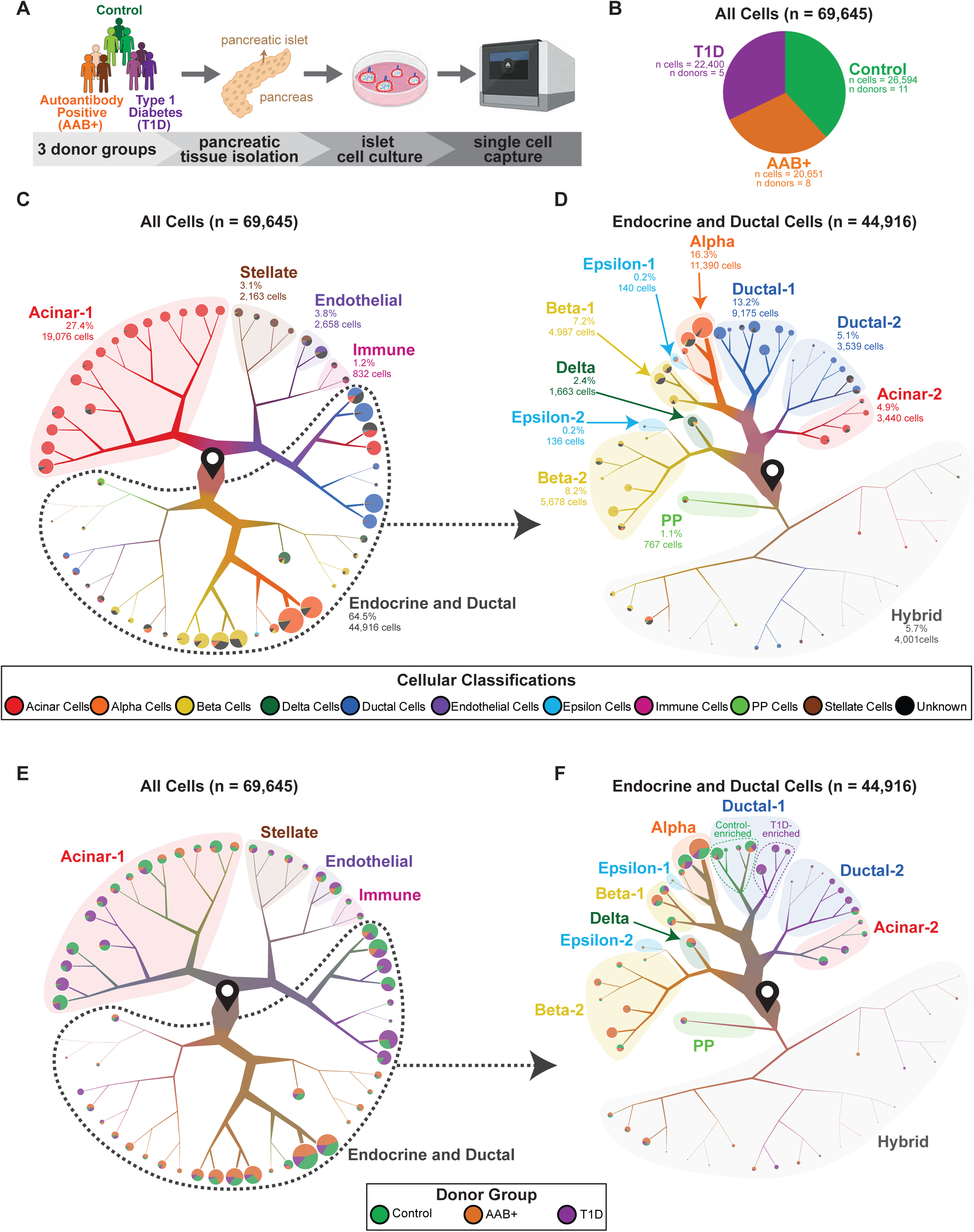
Discernment of human pancreatic cell types using single-cell RNA-seq. (A) The transcriptome of single cells from pancreatic islets of 3 donor types (health Control donors, autoantibody positive (AAB+) donors, and donors with Type 1 diabetes (T1D)) was ascertained using the 10x Genomics platform. (B) Pie chart displaying the proportion of cells comprised by each donor group. (C) TooManyCell dendrogram visualization and clustering of all cells. Cells begin at the start pin symbol, and there are then partitioned based on transcriptional similarities and differences. The color within the branches indicates the proportion of the cells that are classified by the Garnett cellular classification tool. Each bifurcation denotes significant transcriptional differences between the two cell groups. Pie charts at the end of the branches display the breakdown of Garnett cellular classification of cells within that terminal cluster. Highlighting or dotted lines surrounding particular clusters of cells with labels define cell types based on Garnett cellular classifications and canonical gene expression (fig. S4a). Branch thickness and piechart size is proportional to cell number. Branch length is not indicative of any factor, but is merely a means by which to display cells within a defined space. Beta cells (*INS* high), alpha cells (*GCG* high), delta cells (*SST* high), PP cells (*PPY* high), epsilon cells (*GHRL* high), acinar cells (*CPA1* high), ductal cells (*KRT19* high), endothelial cells (*VWF* high), stellate cells (*RSG10* high), and immune cells (*PTPRC,* also known as CD45 or leukocyte common antigen, high). (D) Dendrogram visualization and clustering of ductal and endocrine cells. Highlighting or dotted lines surrounding particular clusters of cells with labels define cell types based on Garnett cellular classifications and canonical gene expression (fig. S4b). (E) Group donor type projected across the dendrogram visualization and clustering of all cells from Figure 1C. Pie charts at the end of the branches display the breakdown donor type within that terminal cluster. (F) Group donor type projected across the dendrogram visualization and clustering of endocrine and ductal cells from Figure 1D. Pie charts at the end of the branches display the breakdown of donor type within that terminal cluster.

Notably, clustering was clearly driven by cell type, as opposed to confounding factors such as autoantibody status, age, BMI, phenotypic group, or other factors (fig. S5, a to k, and fig. S6, a to i). This clustering pattern corroborates the high quality of the resultant data, insinuating the strong biological signal in these scRNA-seq measurements. Additional evidence for the lack of technical noise stems from the observation that cell type clustering was preserved when donors from T1D, AAB+, and Control categories were independently clustered (fig. S7, a to f).

Transdifferentiation of ductal cells to endocrine cells has been reported in models of extreme pancreatic injury (*11–16*). Therefore, we examined the relationship between pancreatic endocrine and ductal cells. First, we subsetted and reanalyzed the endocrine and ductal cells to achieve a more granular clustering (Fig. 1D). Upon reclustering, the major cell types – alpha cells, beta cells, delta cells, epsilon cells, PP cells, ductal cells, and acinar cells – were easily discernible by the cell type classifier and assessment of canonical markers (Fig. 1D and fig. S4, b to d). In instances where there were two transcriptionally distinct canonical cell types (i.e. Beta-1/Beta-2), differential gene expression analysis between populations provided further insights into the underlying molecular differences (table S2 to S5). For example, cells in the Beta-2 cluster expressed higher levels of stress response genes (i.e., *GDF15* and *NPTX2*) when compared to those in the Beta-1 cluster. Notably, comparison of cells in the two Ductal clusters revealed that while cells in the Ductal-1 cluster were enriched for transcription factors (TFs) associated with the endocrine cell fate (i.e., *PDX1* and *NKX6-1*), those in the Ductal-2 cluster only expressed acinar TFs (i.e., *PTF1A* and *GATA4*). These differences in TF gene activation in the two ductal populations were further supported by their position on the dendrogram, and might provide insights into the underlying biology of the acinar-duct-islet axis. For instance, it is plausible that cells in the Ductal-1 cluster might be more prone to transdifferentiation into endocrine cells than those in the Ductal-2 cluster.

Surprisingly, a substantial number of cells (4,001 cells) were not included in these canonical cell type clusters, but formed their own transcriptionally distinct groups in close proximity to one another on the dendrogram. This cluster comprised 5.7% of all profiled cells, with a mixture of cellular classifications and expression of canonical gene markers associated with beta, alpha, ductal, and acinar cells. Therefore, we labeled these cells as ‘Hybrid’ cells (Fig. 1D and fig. S4b). Notably, the gene expressed most highly and consistently in the Hybrid cell population was *INS* (fig. S4b), and a comprehensive examination of the cells comprising this cluster ruled out the possibility of them being potential doublets (fig. S1d and fig. S2 a to e).

Since immune cell-mediated autoimmune destruction of viable pancreatic cells is a pathognomonic feature of T1D, we further examined the intrapancreatic immune cells profiled by scRNA-seq. First, we subsetted and reclustered the cells comprising the ‘Immune’ cluster from the comprehensive tree (Fig. 1C), and found that this population also contained stellate (*RGS5* high) and Schwann (*PLP1* high) cells along with immune cells (*PTPRC*) (fig. S8a and b). Using the Immunological Genome Project (ImmGen) cell type signatures (*17*), we further found antigen-presenting cells (APCs) and macrophages gene signatures, for example *CD68, SPI1, CD14,* and *CD16,* were most frequently upregulated in immune cell subset (fig. S8b and c), suggesting that these cell types comprise the majority of the identified immune cells.

In addition to successful identification of major endocrine and exocrine cell types and pancreatic immune cells, we also observed that the overall proportions of cell types was in accordance with previous work (*5, 18–22*). Each of the major identified cell types comprised of cells from the three donor groups with varying proportions (Fig. 1, E and F, fig. S8, d to e, and fig. S9 a to b). As expected, we found that there was a lower proportion of beta cells in the T1D cohort compared to the AAB+ or Control groups (fig. S9 a to b). Conversely, both acinar and ductal cells comprised a higher portion in the T1D cohort, reflecting the difficulty of isolating high purity islets from T1D donors. Furthermore, within major cell clusters, there were varying degrees of separation based on donor group, which is to be expected due to likely transcriptomic differences among the three donor states (Fig. 1, E to F, and fig. S8a). Notably, Ductal-1 cells clearly separated into distinct T1D-enriched and Control-enriched groups (Fig. 1F). Taken together, our data indicate that transcriptomic differences amongst cell types and not technical biases drive separation of major cellular clades, and the donor state further segregates each cell type.

### AAB+ and T1D endocrine and exocrine cells have common and distinct gene expression programs

Beta cell autoantibodies can be detected in the bloodstream of T1D patients months to years prior to clinical diagnosis. To investigate the molecular events that occur in a possibly prediabetic state, we next compared transcriptomic divergence of AAB+ and T1D cells from Controls (Fig. 2A and tables S6 to S35). For the most part, the degree of overlap between dysregulated genes and pathways in AAB+ and T1D states were cell type-dependent (Fig. 2, B to E, fig. S10, a to b, fig. S11, a to b, fig. S12, a to d, fig. S13, a to d, and fig. S14, a to c). However, some pathways were found to be commonly dysregulated in multiple cell types across T1D and AAB+; including ‘apoptotic signaling’, various protein folding ontologies, various viral-related ontologies, ‘autophagy’, ‘inflammatory pathways’, and ‘stress response’.

**Figure 2:**
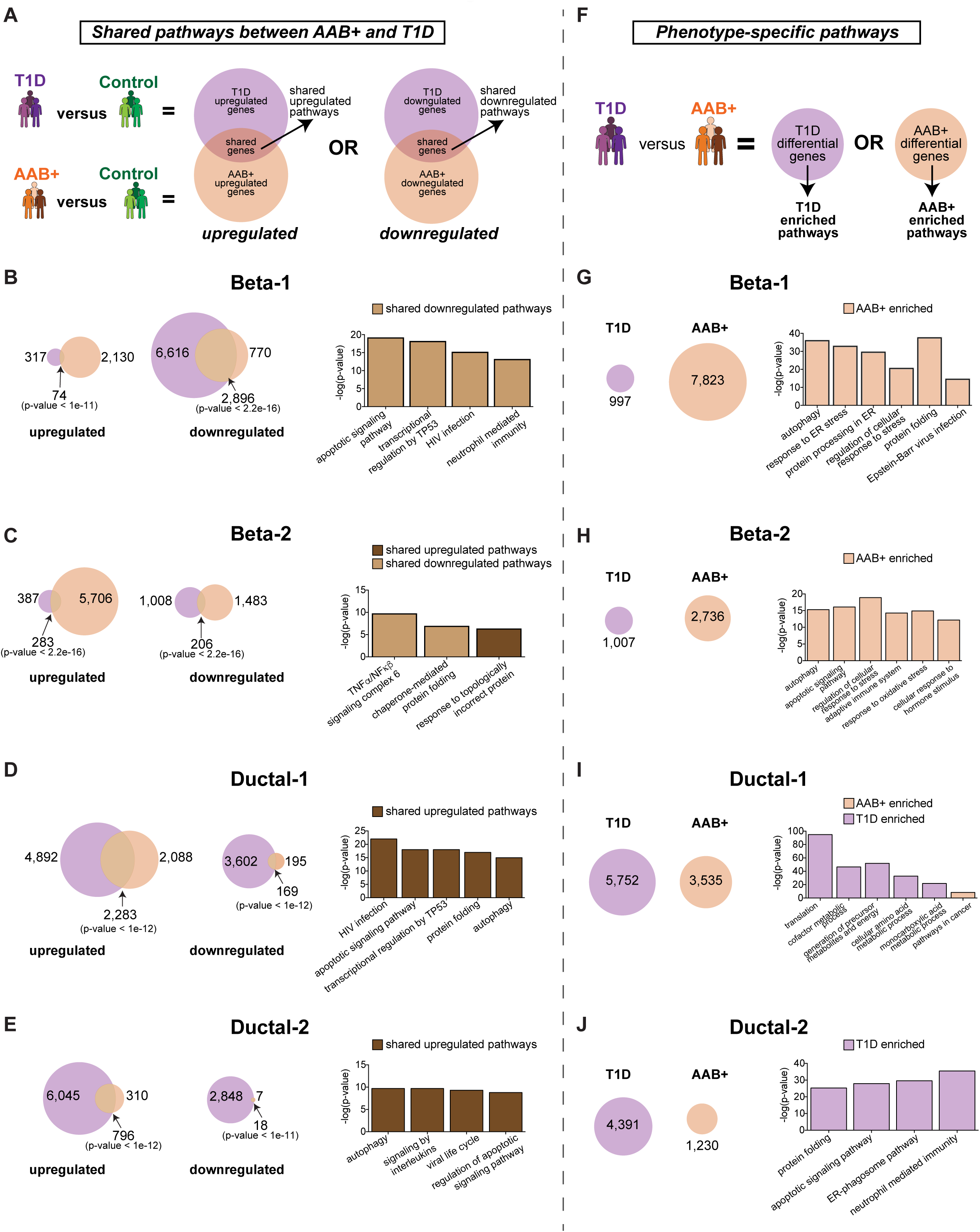
AAB+ and T1D donors have both common and distinct transcriptomic changes in endocrine and exocrine cell types. (A) For each cell type, two pairwise differential comparisons were carried out: (1) T1D versus Control (referred to as ‘T1D upregulated’ (T1D/Control) or ‘T1D downregulated’ (Control/T1D)) and (2) AAB+ versus Control (referred to as ‘AAB+ upregulated’ (AAB+/Control) or ‘AAB+ downregulated’ (Control/AAB+)). T1D upregulated genes were then compared to AAB+ upregulated genes to find commonly upregulated genes, and subsequently commonly upregulated gene ontologies and pathways, across these two donor groups; this exact same approach was carried out for downregulated genes as well. (B-E) (Left) For each cell type, Venn diagrams indicate the numbers of upregulated and downregulated genes, as well as overlapping genes, across the two donor states. (Right) Bar graph displaying notable gene ontologies that are shared across disease states for upregulated and downregulated genes. P-values presented are the results of hypergeometric CDF tests (one-tailed test for overrepresentation). (F) Transcriptional differences between cells from T1D and AAB+ donors were determined by directly comparing T1D to AAB+ cells to generate lists of differentially expressed genes that are enriched in T1D cells or AAB+ cells, and enriched gene ontology pathways were discovered from these differential gene lists. (G-J) (Left) For each cell type, circles indicate the numbers of genes that are ‘T1D enriched’ or ‘AAB enriched’. (Right) Bar graph displaying notable gene ontologies that are enriched for each donor state.

Given the importance of beta cells in T1D pathogenesis, we next examined the transcriptional changes in the two populations of annotated beta cells, Beta-1 and Beta-2. A large number of genes were downregulated in T1D (9,512 genes) and AAB+ (3,666 genes) Beta-1 cells compared to Controls, which frequently overlapped (2,896 genes, 28%; p < 2.2e-16) between the two donor groups (Fig. 2B and fig. S10a). Notable pathways that were frequently downregulated in Beta-1 cells were immune/stress response and apoptosis related (Fig. 2B and fig. S10a). Given that beta cells are destroyed by immune cells in T1D, it is possible that these remaining Beta-1 cells captured in our analysis are able to survive and function by downregulating immune and apoptotic signaling. Notably, these results also suggest that cells from AAB+ donors in this beta cell population employ similar protective molecular mechanisms to enhance survival and function. The Beta-2 cell population displayed a small proportion of genes (283 genes; 4%) that were elevated in both T1D and AAB+ cells when compared to Controls (Fig. 2C, and fig. S10b). Additionally, an even smaller number of genes were downregulated in T1D and AAB+ Beta-2 cells when compared to Controls (Fig. 2C and fig. S10b). Nonetheless, a few gene pathways were found to be commonly dysregulated across both donor groups. Two interrelated pathways dysregulated in both T1D and AAB+ Beta-2 cells suggest a dysregulation of protein folding, an essential function for cellular homeostasis. Additionally, the TNF signaling pathways, which have been implicated as an important regulator of autoimmune processes (*23*), were significantly downregulated across the two donor groups (Fig. 2C and fig. S10b; p-value < 2.2e-16). Together, our differential expression analyses complement earlier studies on the pathways triggering beta cell dysfunction and death, and can further enable the identification of endogenous cellular responses to restore homeostasis at asymptomatic early stages.

Given the clear segregation of ductal cell populations based on the donor group, we next sought to examine the transcriptional changes in the two populations of ductal cells, Ductal-1 and Ductal-2. A large number of genes were upregulated in T1D (7,175 genes) and AAB+ (4,371 gene) Ductal-1 cells when compared to Controls, a significant number of which were common between the two donor groups (Fig. 2D and fig. S11a; 2,283 genes; 25%; p-value < 1e-12). Notable pathways that are upregulated in T1D and AAB+ cells are pathways involved in apoptosis, stress, and immune response (Fig. 2D and fig. S11a). In the Ductal-2 cell population, although a large number of upregulated genes were observed in T1D (6,841 genes), there were not nearly as many upregulated genes in AAB+ cells (1,106 genes) when compared to Controls (Fig. 2E and fig. S11b). Furthermore, in the T1D and AAB+ Ductal-2 cell population, there was a modest but significant overlap between upregulated genes (Fig. 2E and fig. S11b; 11%; p-value < 1e-12). Nevertheless, various gene pathways were found to be significantly upregulated across Ductal-1 cells of both disease states, including apoptotic, autophagy, and immune signaling pathways (Fig. 2, D and E, and fig. S11, a and b). Taken together, these findings suggest that although AAB+ donors maintain normoglycemia, significant transcriptional dysregulation is occurring in AAB+ endocrine and exocrine cells that is highly similar to that in T1D.

Next, we directly compared T1D to AAB+ cells (Fig. 2F and table S36 to S50). For both groups of beta cells, genes associated with apoptosis, stress, and immune response pathways were found to be upregulated in AAB+ cells compared to T1D cells (Fig. 2, G and H, and fig. S10, a and b). For AAB+ Beta-1 cells, it is possible that cells with ‘Epstein-Barr virus infection’ pathway activation are able to survive due to the upregulation of the ‘autophagy pathway’, since this pathway is known to enhance survival over apoptotic or necrotic pathways (*24*); this line of reasoning may also apply to pathways that quell cellular stressors, such as ER stress (Fig. 2, G and H, and fig. S10, a and b). In addition to several commonly upregulated pathways in Beta-1 and Beta-2 AAB+ cells, apoptotic and adaptive immunity signaling were only upregulated in Beta-2 AAB+ cells. These data suggest that this population is undergoing cell death, indicated by the expression of adaptive immune cell genes and *BCL10*. In Ductal cell populations, there was a larger number of upregulated genes in T1D (Fig. 2, I and J, and fig. S11, a and b). More specifically, apoptotic, stress, and immune responses were activated in T1D ductal cells in comparison to AAB+ ductal cells (Fig. 2, I and J, and fig. S11, a and b). Remarkably, interferon alpha and beta pathways, known to be critical in T1D disease pathogenesis (*25–27*), were significantly elevated in T1D ductal cells compared to either Control or AAB+ ductal cells (fig. S14d). While ductal cells have not historically been implicated in T1D, these findings position these exocrine cells in disease pathogenesis.

Taken together, our data indicate that both endocrine and exocrine pancreatic cell types show transcriptional dysregulation in T1D. Remarkably, AAB+ cells exhibit significant transcriptional changes similar to those observed in T1D.

### Single-cell RNA-seq profiling identifies non-canonical, rare Hybrid cells

We next sought to gain a deeper understanding of the rare, non-canonical cell population, which we termed ‘Hybrid’ cells (Fig. 1D). This group consisted of 4,001 cells (5.7% of all cells) with a large fraction of the cells expressing high levels of *INS*, with additional activation of the gene signatures associated with alpha, ductal, and acinar cells (Fig. 1D and fig. S4b). We found that approximately 1.3% of cells from each individual donor (median across all donors) were classified as Hybrid cells, with the exception of one ABB+ donor (AAB+ #3), which was overrepresented in the Hybrid cluster (Fig. 1F, fig. S6d, and fig. S9, a and b). However, the Hybrid cluster was present at a similar frequency even when this AAB+ donor was removed, providing evidence of the presence of this cellular state across multiple donors (fig. S15, a and b).

Detailed interrogation and cellular classifications revealed three distinct subpopulations across the Hybrid cluster (Fig. 3, A and B). The ‘Ductal-like Hybrids’ subpopulation expressed ductal cell signature genes; the ‘Endocrine-like Hybrids’ subpopulation exhibited alpha-cell gene signature. Finally, the third subpopulation of Hybrid cells, called ‘Acinar-like Hybrids’ expressed acinar cell signature genes (Fig. 3. A and B). In addition to expressing beta cell marker *INS*, many Hybrid cells across all three subpopulations co-expressed another marker gene indicative of two independent canonical cell types (Fig. 3C, and fig. S15c).

**Figure 3:**
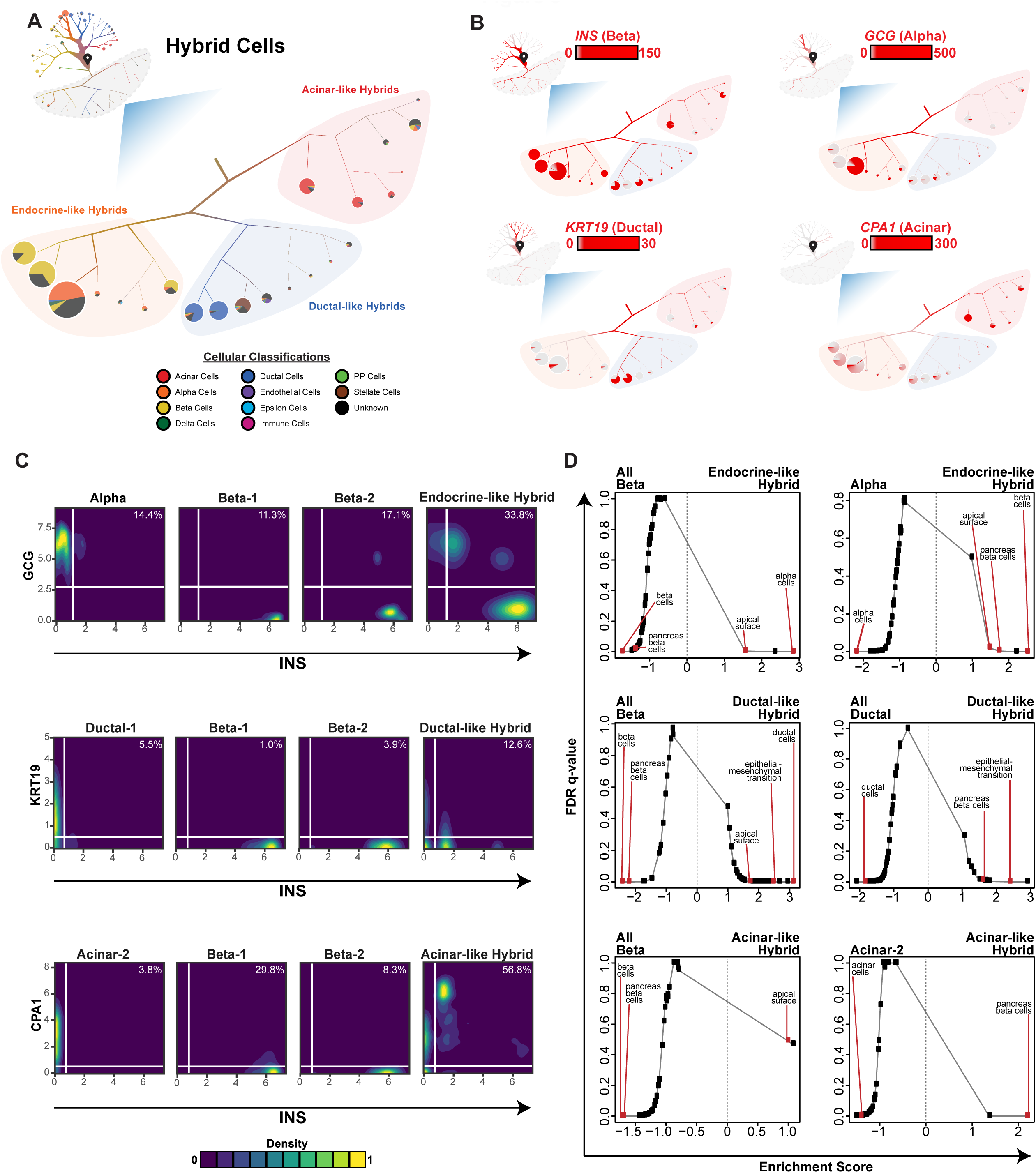
Single cell RNA-seq profiling identifies non-canonical, rare Hybrid cells. (A) (Inset) Dendrogram visualization and clustering of ductal and endocrine cells from Figure 1D. Highlighted in grey are Hybrid cells. (Main) Magnified view of the Hybrid cell population from the ductal and endocrine dendrogram. The color within the branches indicates the proportion of the cells that are classified by the Garnett cellular classifier. Cells begin at the start pin symbol, and from there are partitioned based on transcriptional similarities and differences. Pie charts at the end of the branches display the breakdown of Garnett cellular classification of cells within that terminal cluster. Branch thickness and pie chart size is proportional to cell number. Branch length is not indicative of any factor, but is merely a means by which to display cells within a defined space. (B) Dendrogram visualization of the expression of each canonical gene marker for each major cell type across the dendrograms from A. Scale bars represent normalized transcript numbers. (C) Co-expression analysis of *INS* and the relevant gene indicative of each subpopulation of Hybrid cells (*KRT19* for Ductal-like Hybrids, *GCG* for Endocrine-like Hybrids, and *CPA1* for Acinar-like Hybrids) at the single-cell level across each Hybrid subpopulation and related canonical cell types. For each gene, values represent normalized transcript numbers. Percentages in the upper right-hand corner indicate the percentage of co-expressing cells. (D) GSEA analysis plots of FDR q-value vs Normalized Enrichment Score. Each Hybrid cell subpopulation was compared to relevant canonical cell populations to determine differentially enriched gene sets. Labels in red demarcated signatures of interest.

To gain further insight into the molecular nature of Hybrid cells, we next compared each Hybrid subpopulation to relevant canonical cell populations to determine their transcriptional divergence (tables S51 to 57) and convergence with respect to gene signatures of common pancreatic cell types (*22, 28*). To this end, we observed the enrichment of the alpha cell signature in Endocrine-like Hybrids, the ductal cell signature in Ductal-like Hybrids, and the acinar cell signature in Acinar-like Hybrids. Conversely, when each of the Hybrid subpopulations was compared to relevant canonical cell types, the beta cell signature was found to be significantly associated with the Hybrid subpopulations (Fig. 3D).

We next investigated the expression of key endocrine, ductal, and acinar transcription factors (TFs) (*29*) in Hybrid cells (tables S51 to S57). Compared to exocrine cells, Hybrid cells were found to have an enrichment of endocrine TF genes (positive log_2_ fold change) (fig. S15d). Conversely, compared to endocrine cells, Hybrid cells were found to have an enrichment of exocrine TF genes (fig. S15d). Furthermore, pseudotime analysis (*30*) of Hybrid cells amongst either exocrine-related or endocrine-related cells demonstrated that the Hybrid cell population is a transcriptionally distinct cellular state with an intermediary mixture of canonical cell types (fig. S16, a to f).

Finally, we assessed standard hallmark gene signatures to determine whether general pathways are enriched in Hybrid cells in comparison to canonical cell types. Across multiple comparisons, Hybrid cells were found to be enriched with the ‘epithelial-mesenchymal transition’ and ‘apical surface’ signatures (Fig. 3D and fig. S15e). Notably, the ‘epithelial-mesenchymal transition’ signature has been implicated in the conversion of ductal cells to endocrine cells (*31, 32*). Taken together, our findings suggest that Hybrid cells are a transcriptionally distinct population of cells across all donors with disparate, yet intermediary, transcriptional signatures of two canonical cell types.

### Proteomic validation of Hybrid cells by mass cytometry

Although gene expression measurement by droplet-based scRNA-seq is unbiased, doublets are observed ∼1% of the time. We used strict criteria and computationally removed potential doublets from scRNA-seq analysis using two independent methods (see Methods). Nevertheless, to address the possibility of doublet artifact in detection of Hybrid cells, we first employed cytometry by time-of-flight (CyTOF) (*33*). Considering the relatively low frequency of Hybrid cells predicted by scRNA-seq (∼5%), we took an integrative approach and combined more than 6 million live single cells from 12 donors, which have also been profiled by scRNA-seq (4 Control, 4 AAB+, and 4 T1D donors). To ascertain if the Hybrid population exists using these deep phenotyping data, we scaled our analytical strategy to millions of cells, and clustered them by expression levels of 33 proteins (table S58). Despite the inherent variability of antibody staining across donors that limited the integrative analysis of previous studies (*34*), we were able to annotate major cell types based on the expression of canonical proteins (fig. S17, a to e).

Remarkably, our strategy identified a cluster of rare cells representing the Ductal-like Hybrid cells previously identified in scRNA-seq, which are cytokeratin/C-peptide double-positive cells (Fig. 4A). The median frequency of Hybrid cells measured by CyTOF across all donors was around 2.7%, similar to the scRNA-seq results, and cells from Control donors constituted a significantly larger proportion of the Hybrid population (Fig. 4, B and C; p-value < 2.2e-16). Finally, the presence of cytokeratin/C-peptide Hybrid cell population was confirmed by two-parameter analysis for donors predicted by our integrative analysis to have the highest and lowest frequency of Hybrid cells (Fig. 4D and fig. S17o).

**Figure 4:**
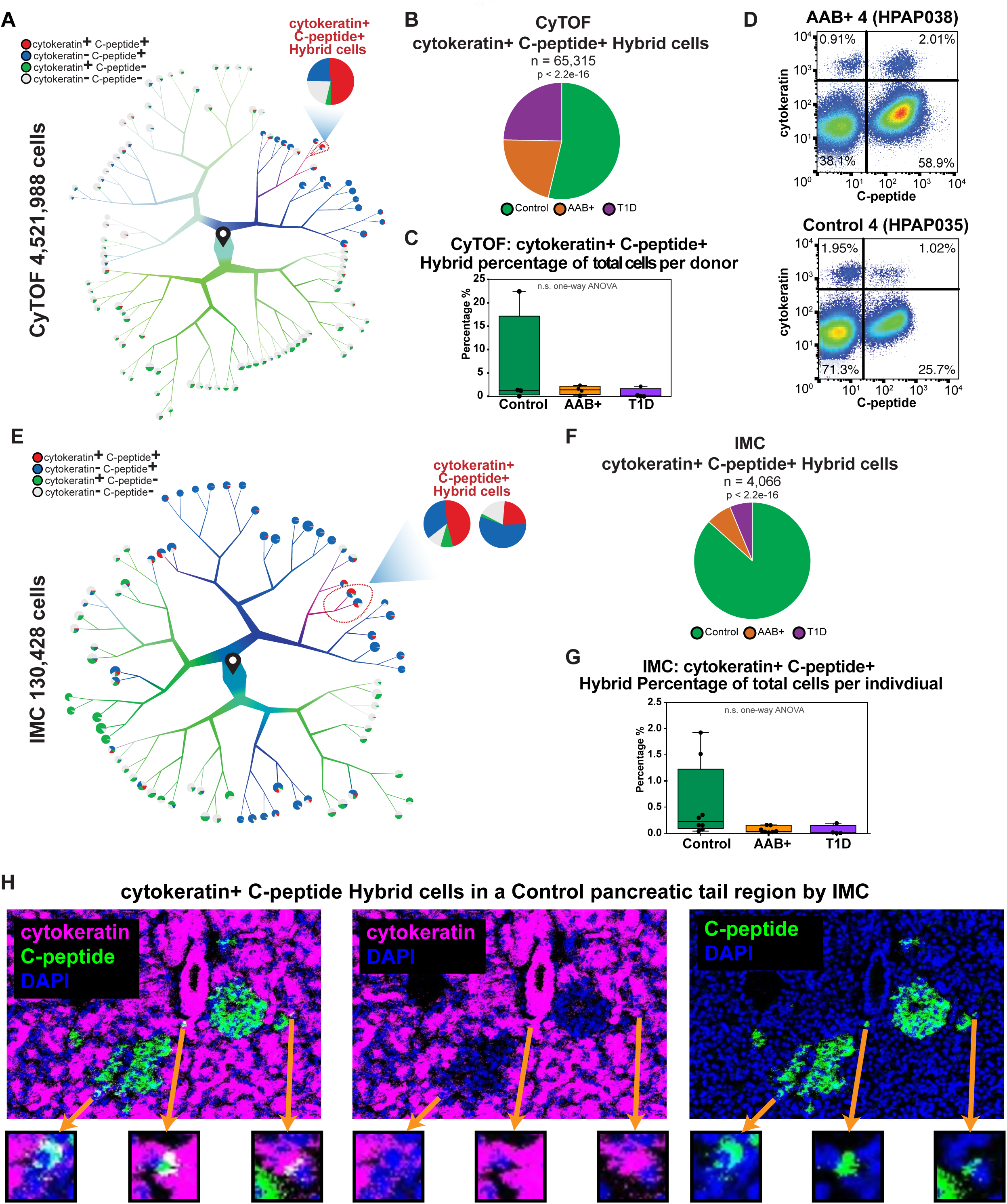
Flow cytometry by time-of-flight (CyTOF) and imaging mass cytometry (IMC) validates the presence of Hybrid cells. (A) Dendrogram visualization of co-expression of cytokeratin and C-peptide proteins in individual cells analyzed with flow cytometry by time-of-flight (CyTOF). Cells begin at the start pin symbol, and from there are partitioned based on similarities and differences in protein markers. (B) Pie chart displaying cytokeratin+ C-peptide+ Hybrid cells and the relative proportions of each donor group from the CyTOF data. P-values are the result of the Chi-squared test. (C) Box plots displaying Hybrid cell percentage of total cells per individual across donor groups from the CyTOF data. (D) Two parameter CyTOF analysis of cytokeratin and C-peptide expression in single cells from AAB+ donor #4 (HPAP038) and Control donor #4 (HPAP035). (E) Dendrogram visualization of co-expression of cytokeratin and C-peptide proteins in individual cells analyzed by imaging mass cytometry (IMC). Cells begin at the start pin symbol, and from there are partitioned based on similarities and differences in protein markers. (F) Pie chart displaying cytokeratin+ C-peptide+ Hybrid cells and the relative proportions of each donor group from the IMC data. P-values are the result of the Chi-squared test. (G)Box plots displaying Hybrid cell percentage of total cells per individual across donor groups from the IMC data. (H) Representative region of interest (ROI) from the tail pancreatic region of a Control donor displaying three cytokeratin+C-peptide+ Hybrid cells.

### Imaging mass cytometry in intact tissues corroborates the presence of Hybrid cells

Having confirmed the existence of Hybrid cells by two orthogonal data modalities, we next sought to study the anatomical-spatial features of Hybrid cells in pancreatic tissues using imaging mass cytometry (IMC) (*5*). Although CyTOF and scRNA-seq rely on the profiling of dissociated cells, IMC removes this confounding effect by analyzing tissues fixed directly from the native human pancreas. We again amended our analytical pipeline to harness the expression levels of 35 proteins quantified by IMC in more than 1 million cells across 147 tissue slides from 19 donors, including 11 individuals not previously assessed by scRNA-seq or CyTOF for an independent validation of our findings (table S59).

IMC analysis not only discerned Ductal-like Hybrid cells amongst ductal and endocrine cells (Fig. 4E and fig. S17, f to j), but also quantified the frequency of Hybrid cells across donor groups (Fig. 4, F and G). Considering that pancreatic tissue sections contain much larger cellular heterogeneity than islet cultures, we found that Hybrid cells accounted for ∼0.4% of all cells across donors (Fig. 4, F and G). Despite normalizing the proportion of Hybrid cells by the number of cells per donor, we found that Controls accounted for significant majority of the Hybrid population (Fig. 4 F-G; p-value < 2.2e-16). We also found that the frequency of immune cells did not correlate with the frequency of Hybrid cells across donor groups (Fig. 4, A, B, C, E, F, and G, and fig. S17, a, d, e, f, i, j, k, and l).

Next, we leveraged the IMC map of spatial architecture of human islets and determined the location of Hybrid cells within the pancreas and interrogated their cellular neighborhood. We found Hybrid cells most frequently located in close proximity to beta and alpha cells, although some Hybrid cells were located near ductal cells (fig. S17, m and n; p-value < 1e-2). Interestingly, the frequency of Hybrid cells was highest in the tail region of the pancreas (Fig. 4H, fig. S17p). Taken together, our multimodal, single-cell profiling approach using scRNA-seq, CyTOF, and IMC enabled the identification and substantiation of non-canonical, rare Hybrid cells. Since the enrichment of Hybrid cells in a particular donor group was not consistent across three technologies, we conclude that Hybrid cells represent a general feature of pancreatic tissues.

### MHC Class II expression is enriched in T1D ductal cells

The major genetic susceptibility determinants of T1D have been mapped to the MHC Class II genes (*35, 36*). We therefore sought to determine which cell types or donor states disproportionately express genes in this pathway. Using our scRNA-seq data, we found that genes associated with MHC Class II activity were enriched in Immune, Endothelial, Ductal, and Ductal-like Hybrid clusters (fig. S18, a to d). The lack of enrichment of the immune cell marker *PTPRC* or other genes associated with immune cells across the endocrine and ductal dendrogram supports that this finding is not due to immune cell contamination (fig. S4,a and b, fig. S18g). We next evaluated the expression of *HLA-DPB1*, an MHC class II gene associated with T1D risk, and *KRT19*, a ductal cell marker, across ductal and endocrine cell types. We identified five clusters with high *HLA-DPB1* and high *KRT19* expression, which accounted for 10.9% of all cells (7,588 cells) (Fig. 5, A and B, fig. S18, e and f). Strikingly, cells from T1D donors disproportionately contributed to the population of MHC Class II-expressing ductal cells (Fig. 5C; p-value < 2.2e-16). This observation is not due to sampling issues pertaining to the difficulty of isolating high purity islets from T1D donors. The Ductal-1 cell population consists of very similar numbers of Control and T1D donor ductal cells (4,217 and 4,154 cells, respectively); however, there is a marked difference in the percentage of Control versus T1D MHC class II-expressing Ductal-1 cells, at 35% and 91%, respectively (Fig. 5C; p-value < 2.2e-16).

**Figure 5:**
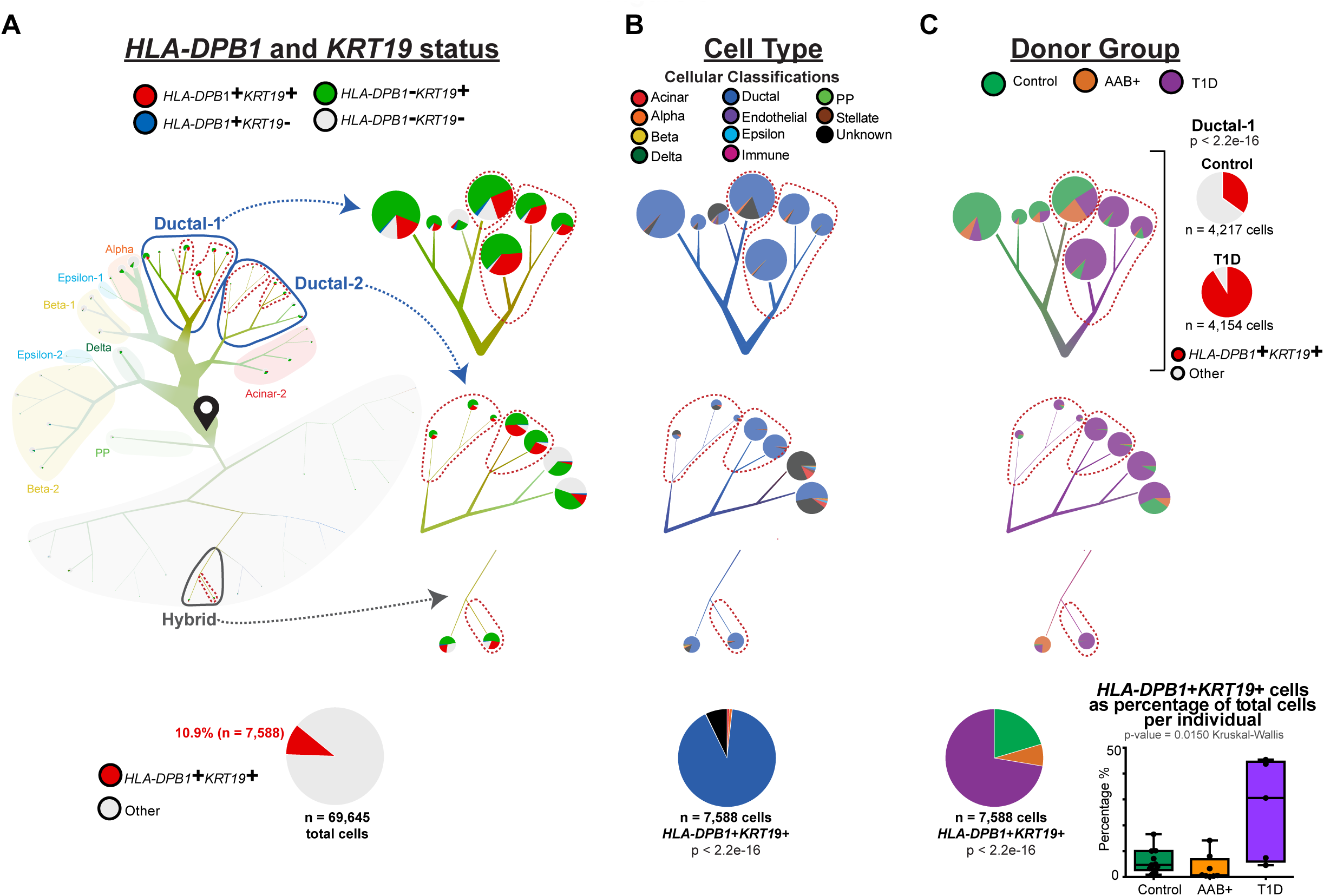
Single cell RNA-seq profiling enables the identification of MHC Class II expressing ductal cells enriched in T1D. (A) (Left Top) Dendrogram visualization of co-expression of *HLA-DPB1* and *KRT19* gene transcripts in individual cells by scRNA-seq across the ductal and endocrine dendrogram from Figure 1D. (Left Bottom) Pie chart demonstrating *HLA-DPB1+KRT19+* cells as percentage of total cells. (Right) Magnified view of the clusters of cells with high percentage (25% or greater) of *HLA-DPB1+KRT19+* cells with *HLA-DPB1* and *KRT19* status displayed across these clusters (outlined in red dashed lines) and neighboring clusters of cells. Cells begin at the start pin symbol, and from there are partitioned based on similarities and differences in gene expression. (B) (Top) Dendrogram visualization of cellular classification status across the magnified clusters of cells with high percentage (25% or greater) of *HLA-DPB1+KRT19+* cells (outlined in red dash lines) and neighboring clusters of cells. (Bottom) Pie chart displaying the relative proportion of cellular classification status of *HLA-DPB1+KRT19+* cells. P-value presented is the result of the Chi-squared test. (C)(Top Left) Dendrogram visualization of donor group across the magnified clusters of cells with high percentage (25% or greater) of *HLA-DPB1+KRT19+* cells (outlined in red) and neighboring clusters of cells. (Top Right) Pie chart displaying the relative proportion of *HLA-DPB1+KRT19+* cells in Control (top) or T1D (bottom) Ductal-1 cells. P-value presented is the result of the Fisher exact test. (Bottom Left) Pie chart displaying the relative proportion of donor group of *HLA-DPB1+KRT19+* cells. P-value presented is the result of the Chi-squared test. (Bottom Right) Box plots displaying the *HLA-DPB1*+*KRT19*+ cell percentage of total cells per individual across donor groups.

### Multimodal confirmation of MHC Class II-expressing ductal cells in T1D

Based on our results from single-cell transcriptomic profiling of the human pancreas (Fig. 5), we surmised that MHC Class II molecules on ductal cells might be involved in antigen presentation in T1D. To validate this result, we employed two high-throughput technologies, CyTOF and IMC, in addition to immunofluorescence experiments in native pancreatic tissues. Using CyTOF, we identified a population of ductal cells expressing HLA-DR, an MHC Class II protein (Fig. 6A). Notably, we found that cells from T1D donors constituted the largest percentage of this cluster, in agreement with the findings from scRNA-seq (Fig. 6B; p-value < 2.2e-16). Furthermore, HLA-DR-expressing ductal cells made up a large percentage of total cells across individual T1D donors (Fig. 6C). A two parameter (cytokeratin and HLA-DR) analysis on all single cells analyzed by CyTOF further confirmed the presence of this double-positive population across multiple donors (Fig. 6D and fig. S19, a and b). Notably, these ductal cells did not express CD45, the hallmark of leukocytes (Fig. 6E).

**Figure 6:**
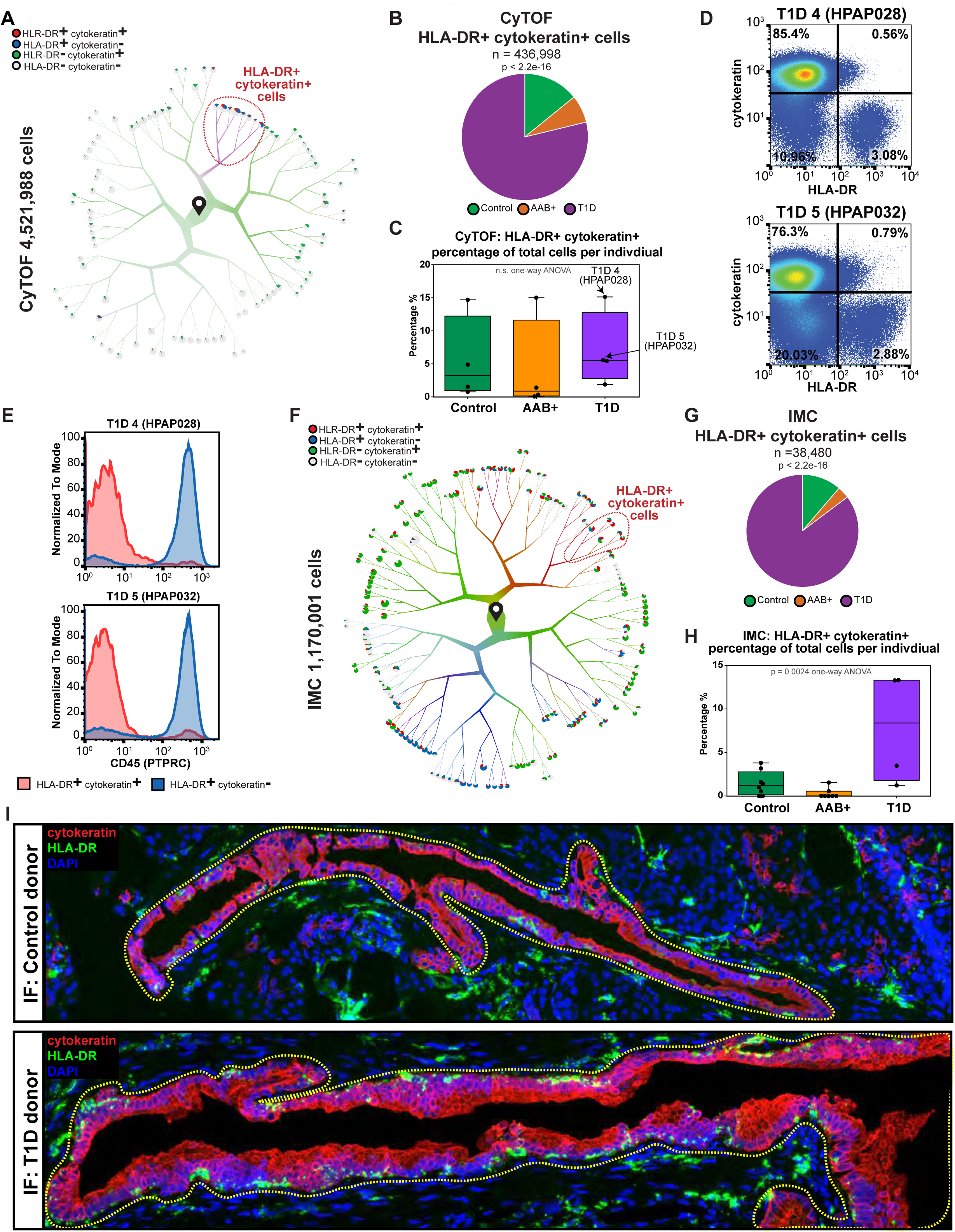
Three single-cell resolution protein-based approaches corroborate the MHC Class II-expressing ductal cells in T1D. (A) Dendrogram visualization of co-expression of HLA-DR and cytokeratin proteins in single cells analyzed with flow cytometry by time-of-flight (CyTOF). Cells begin at the start pin symbol, and from there are partitioned based on similarities and differences in protein levels. (B) Pie chart displaying HLA-DR+ cytokeratin+ cells and the relative proportions of each donor group from the CyTOF data. P-value presented is the result of the Chi-squared test. (C) Box plots displaying HLA-DR+ cytokeratin+ cell percentage of total cells per individual across donor groups from the CyTOF data. (D) Two parameter CyTOF analysis of HLA-DR and cytokeratin protein expression in single cells from T1D donor #4 (HPAP028) and T1D donor #5 (HPAP032). (E) CD45 (PTPRC) expression levels in HLA-DR+ cytokeratin+ and HLA-DR+ cytokeratin-single cells. (F) Dendrogram visualization of co-expression of HLA-DR and cytokeratin proteins in single cells analyzed by imaging mass cytometry (IMC). Cells begin at the start pin symbol, and from there are partitioned based on similarities and differences in protein levels. (G)Pie chart displaying HLA-DR+ cytokeratin+ cells and the relative proportions of each donor group from the IMC data. P-value presented is the result of the Chi-squared test. (H) Box plots displaying HLA-DR+ cytokeratin+ cell percentage of total cells per individual across donor groups from the IMC data. (I) Representative region of interest (ROI) from the pancreas of a T1D donor (top) or Control donor (bottom) displaying HLA-DR+ cytokeratin+ cells via immunohistochemistry (IHC) followed by confocal microscopy.

To comprehensively study the existence of MHC Class II-expressing ductal cells in pancreatic tissues independent of islet cultures, we exploited the quantitative analysis of IMC data. By analyzing 147 tissue slides, we confirmed that MHC Class II-expressing ductal cells were predominately present in T1D donors (Fig. 6, F to H, fig. S20). MHC Class II-expressing ductal cells were located primarily in the tail and body regions of the pancreas (fig. S19c). Remarkably, the frequency of immune cells highly correlated with the frequency of MHC Class II expressing ductal cells, given that the T1D donor group had the highest percentage of both immune cells and HLA-DR-expressing ductal cells (fig S19, d and e). Finally, immunofluorescence staining (IF), followed by confocal microscopy verified the existence of MHC Class II-expressing cells in the pancreatic tissues of a Control and a T1D donor. We identified MHC Class II-expressing ductal cells in both donors; however, there was a more pronounced enrichment of MHC Class II-expressing ductal cells in the T1D pancreas (Fig. 6I). Cellular neighborhood analysis in pancreatic tissues further unveiled that HLA-DR-expressing ductal cells were surrounded by immune cells, in particular CD4^+^ immune cells and CD11b^+^ dendritic cells (fig. S19, f and g; p-value < 1e-2). Together, our multi-modal single-cell measurements from transcriptomics to spatial proteomics in ductal cells provide novel insights into pancreas biology and suggest that ductal cells could be involved in the initial pathogenic events of T1D by functioning as antigen-presenting cells.

## Discussion

Here, we employed scRNA-seq as a molecular microscope to investigate cellular diversity in pancreatic islets of T1D, AAb+, and non-diabetic human organ donors, comprehensively analyzing over 80,000 cells. Importantly, a web portal of the scRNA-seq data presented here is publicly available for exploration (*37*). We further leveraged high-throughput proteomic assays, including mass cytometry (CyTOF) and multiplexed histological analyses by IMC, to corroborate transcriptomic-based findings.

We found that AAb+ donors exhibit similar transcriptional changes to T1D donors in various endocrine and exocrine cells, despite these donors retaining normoglycemia. This observation suggests that even though AAb+ donors are clinically normal, various cell types of AAb+ donors have a molecular phenotype similar to those of T1D donors. Although it is impossible to discern at present whether these transcriptional changes are contributing to or are byproducts of disease pathogenesis, the mere discovery of molecular phenotypic changes in these cells in AAb+ individuals is novel and advances our understanding of early pancreatic perturbations occurring in T1D.

We identified a rare population of Hybrid cells, with expression signatures of beta, ductal, alpha, and acinar cells. Similar mixed signature cell types have been reported previously using single-cell transcriptomics (*22, 38*). Although the human fetal pancreas has been known to contain multihormonal cells (*39, 40*), as well as cells with dual endocrine-exocrine signatures (*22, 38*), these types of cells have only recently been documented in the adult stage in mice and humans (*22, 38, 41-43*). It is possible that these Hybrid cells in adulthood are multihormonal cells retained from early development or are formed when monohormonal cells differentiate into another cell type during adulthood. Most recently, it has been suggested that multipotent progenitor-like cells in humans defined as PDX1^+^/ALK3^+^/CAII^−^ cells can differentiate into all pancreatic lineages including endocrine cells (*44*). While cells annotated as Hybrid cells in our study did not show high expression levels of *ALK3* (*BMPR1A*) (fig. S18h), it is likely that large scale investigations of exocrine cells will shed more light onto the cellular heterogeneity and regenerative potential of this compartment.

The most striking finding arising from our study is that cells of the exocrine compartment show transcriptional and gene ontological changes in the T1D disease setting. Ductal cells from T1D donors, in a sharp contrast with non-diabetic or AAb+ donors, expressed high levels of MHC Class II and interferon pathways and were surrounded by immune cells. Although our study represents the first report of ductal cells expressing MHC Class II proteins in the T1D context, this finding is in accordance with previous literature documenting an elevation of immune cells in the exocrine pancreas of T1D donors (*5, 27, 45*) and regulation of MHC Class II genes by the interferon signaling pathway (*46*). Recent reports support a role for epithelial cells as facultative, non-professional antigen presenting cells in the gut and lung (*47*), and expression of MHC Class II proteins in non-lymphoid cells in the pancreas (*48*). Notably, two HLA genes with high expression in ductal cells, *HLA-DRB1* and *HLA-DPB1*, are major T1D susceptibility genes (*35*). Together, our study unmasks exocrine ductal cells as potential pathogenic cells in T1D and suggest a new cellular context for examining the role of genetics in T1D.

## Acknowledgments

We thank our colleagues for helpful discussions, particularly: Aditi Chandra, Eline Luning Prak, Ben Stanger, Michael Silverman, Gregory Beatty, Ken Zaret, Mitch Lazar, and E. John Wherry. We thank Andrei Georgescu for confocal microscopy and the University of Pennsylvania Diabetes Research Center (DRC) for the use of the Functional Genomics Core (P30-DK19525). This work was supported by NIH grants UC4 DK112217 (to A.N., K.K., M.B., J.M., M.F., and G.V.), R01CA230800 and Susan G. Komen CCR185472448 (to R.B.F.), and R01HL145754, U01DK127768, the Burroughs Wellcome Fund, the Chan Zuckerberg Initiative, the Penn Epigenetics and the Sloan Foundation awards to G.V.

## Materials and Methods

### Experimental model and subject details

Pancreatic islets were procured by the HPAP consortium under the Human Islet Research Network (https://hirnetwork.org/) with approval from the University of Florida Institutional Review Board (IRB # 201600029) and the United Network for Organ Sharing (UNOS). A legal representative for each donor provided informed consent prior to organ retrieval. For T1D diagnosis, medical charts were reviewed and C-peptide levels were measured in accordance with the American Diabetes Association guidelines (American Diabetes Association 2009). All donors were screened for autoantibodies prior to organ harvest, and AAb positivity was confirmed again post tissue processing and islet isolation.

Organs were recovered and processed as previously described (*7*). Table 1 summarizes donor information. Pancreatic islets were cultured and dissociated into single cells as previously described (*22*). Total dissociated cells were used for single cell capture for each of the donors, except AAB+ donor #1 (HPAP019), which was enriched for beta cells as previously reported prior to capture (*49*).

### scRNA-seq islet capture, sequencing, and processing

The Single Cell 3’ Reagent Kit v2 or v3 was used for generating scRNA-seq data. 3,000 cells were targeted for recovery per donor. All libraries were validated for quality and size distribution using a BioAnalyzer 2100 (Agilent) and quantified using Kapa (Illumina). For samples prepared using ‘The Single Cell 3’ Reagent Kit v2’, the following chemistry was performed on an Illumina HiSeq4000: Read 1: 26 cycles, i7 Index: 8 cycles, i5 index: 0 cycles, and Read 2: 98 cycles. For samples prepared using ‘The Single Cell 3’ Reagent Kit v3’, the following chemistry was performed on an Illumina HiSeq 4000: Read 1: 28 cycles, i7 Index: 8 cycles, i5 index: 0 cycles, and Read 2: 91 cycles. Cell Ranger (10x Genomics; v3.0.1) was used for bcl2fastq conversion, aligning (using the hg38 reference genome), filtering, counting, cell calling, and aggregating (--normalize=none) (fig. S1, a and b).

### scRNA-seq clustering, doublet removal, & cell type classification

Seurat v3.1.5 (*19, 50*) was used for filtering, UMAP generation, and initial clustering. Genes expressed in at least 3 cells were kept, as were cells with at least 200 genes. nFeature, nCount, percent.mt, nFeature vs nCount, and percent.mt vs nCount plots were generated to ascertain the lenient filtering criteria of 200 > nFeature < 8,750, percent.mt < 25, and nCount <125,000. Data was then log normalized, and the top 2,000 variable genes were detected using the “vst” selection method. The data was then linearly transformed (“scaled”), meaning that for each gene, the mean expression across cells is 0 and the variance across cells is 1. Principle component analysis (PCA) was then carried out on the scaled data, using the 2,000 variable genes as input. We employed two approaches to determine the dimensionality of the data, i.e. how many principal components to choose when clustering: (1) a Jackstraw-inspired resampling test that compares the distribution of p-values of each principle component (PC) against a null distribution and (2) an elbow plot that displays the standard deviation explained by each principal component. Based on these two approaches, 17 PCs with a resolution of 1.2 were used to cluster the cells, and non-linear dimensionality reduction (UMAP) was used with 17 PCs to visualize the dataset (fig. S1, c and e).

Two independent methods were used to detect and remove doublets. DoubletFinder v2.0 (*51*) was used to demarcate potential doublets in the data as previously described, with the following details: 17 PCs were used for pK identification (no ground-truth) and the following parameters were used when running doubletFinder_v3: PCs = 17, pN = 0.25, pK = 0.0725, nExp = nExp_poi, reuse.pANN = FALSE, and sct = FALSE (fig. S1d). Scrublet v0.2.1 (*18*) was also used to demarcate potential doublets (fig. S2a). Given that a small percentage of cells were demarcated as doublets by both methods, we removed all cells that were flagged as doublet by both or either approach, leading to the removal of 3,770 cells (fig. S2, b and c).

Following doublet removal, the raw data for the remaining 73,235 cells were filtered using the following criteria, which resulted in 69,645 cells remaining: 200 > nFeature < 8,750, percent.mt < 25, and nCount <100,000 (fig. S2c and fig. S3a). The data were log normalized, the top 2,000 variable genes were detected, the data underwent linear transformation, and PCA was carried out, as described above. Both the Jackstraw-inspired resampling test and an elbow plot of standard deviation explained by each principal component were used to determine the optimal dimensionality of the data, as described above. Based on these two approaches, 26 PCs with a resolution of 1.2 was used to cluster the cells, and UMAP was used with 26 PCs to visualize the 49 clusters detected (fig.S3, b and c).

Garnett was used for initial cell classification as previously described (*9*). In brief, a cell type marker file (table S60) with 17 different cell types was compiled using various resources (*18–22, 28*), and this marker file was checked for specificity using the “check_markers” function in Garnett by checking the ambiguity score and the relative number of cells for each cell type. A classifier was then trained using the marker file, with “num_unknown” set to 500, and this classifier was then used to classify cells and cell type assignments were extended to nearby cells, “clustering-extended type” (Louvain clustering) (fig. S3d). Upon inspection of cluster purity using canonical gene markers of the major pancreatic cell types, we found that the abundant and transcriptionally distinct cell types form generally distinct and unique clusters: beta cells (*INS* high), alpha cells (*GCG* high), acinar cells (*CPA1* high), ductal cells (*KRT19* high), endothelial cells (*VWF* high) stellate cells (*RSG10* high), and immune cells (*PTPRC,* also known as CD45 or leukocyte common antigen, high) (fig. S3e). In contrast, the rarer and/or less transcriptionally distinct cell types did not clearly segregate, namely delta cells (*SST* high), PP cells (*PPY* high), and epsilon cells (*GHRL* high) (fig. S3e).

To overcome the apparent limitation in grouping the major canonical cell types, we employed the analytical workflow termed ‘TooManyCells’ (*10*), which implements an efficient divisive hierarchical spectral clustering approach along with tree visualizations to preserve the relationships among cell clusters at varying resolutions. In conjunction with ‘TooManyCells’ clustering, we invoked the cellular classifier Garnett (*9*), which annotates cell types by training a regression-based classifier from user-provided cell type signatures (table S60; (*34*)). Briefly, for the clustering of all cells, the raw data from the 69,645 cells were normalized by total count and gene normalization by median count (TotalMedNorm) followed by term frequency-inverse document frequency (tf-idf) for clustering. For visualization of the comprehensive clustering, the dendrogram was first pruned using the TooManyCells flags ‘--min-distance-search “15”’ and “--smart-cutoff “15”’, followed by pruning using the flag ‘--max-step 6’. TooManyCells enabled distinct clustering of all major known cell types associated with pancreatic islets, as confirmed by Garnett cell labels and the inspection of canonical gene markers for each cell type (Fig. 1C and fig. S4a).

The raw data from different cell types were then subsetted from the comprehensive clustering in Figure 1C in order to cluster cells on a cell-type basis. For the clustering of ductal/endocrine cells, data from the ductal/endocrine cell clusters from the comprehensive tree were subsetted and normalized by TotalMedNorm followed by term tf-idf. For visualization of the ductal/endocrine tree, the dendrogram was first pruned using the TooManyCells flags ‘--min-distance-search “7”’ and “--smart-cutoff “7”’ followed by pruning using the flag ‘--max-step 7’. For the clustering of immune cells, data from the immune cell cluster from the comprehensive tree were subsetted and normalized by TotalMedNorm followed by tf-idf. For visualization of the immune tree, the dendrogram was first pruned using the TooManyCells flag ‘--max-step 4’. When individual genes were painted across any of the dendrograms, ‘TotalMedNorm’ was employed to normalize gene expression.

### Differential Gene Expression, GSEA analysis, and Metascape analysis

Differential genes were found using edgeR through TooManyCells with the – normalization “NoneNorm” to invoke edgeR single cell preprocessing, including normalization and filtering. For Metascape analysis (http://metascape.org/gp/index.html#/main/step1; (*52*)), less than or equal to 3,000 differential genes (FDR< 0.05 and fold change (FC) > 0.1) were subjected to analysis. The top 20 clusters are displayed and a stringent cut-off of 1e-6 was applied to determine significant gene ontology pathways. For gene-set-enrichment-analysis (GSEA) analysis, GSEA Preranked (4.0.1) (*53*) was run on a pre-ranked gene list using either user-provided pancreatic gene expression sets (*22, 28*) or standard hallmark gene signatures provided by the Molecular Signatures Database (MSigDB) (*54*).

### Hybrid cell co-expression, DE analysis and heatmaps

With the differentially expressed genes (FDR< 0.05 and fold change (FC) > 0.1) between every two sample groups, we calculated the shared and unique genes in each cell type, and visualized it with Venn diagrams. The expression levels of the genes in each cell of the three groups were extracted from the median normalized count matrix. Then we aggregate the expression levels in each group by taking the average value of the normalized counts. The mean expression values of the three groups were further normalized by the total expression level of that gene. We visualized the normalized expression level of differential genes with heatmaps.

To examine the co-expression of signature genes of some cell types, we normalized the median normalized matrix with log2(N +1). Then we selected the matrix of selected cell types by marker genes. The distribution of the cells from selected cell types by expression level of two marker genes were shown with geom_density_2d_filled() in ggplot2 package of R.

### Transcription Factor analysis

A list of human transcription factors was downloaded (*55*), and the expression of key transcription factors (TFs) of endocrine, ductal, and acinar cell fates (*29*) were assessed in Hybrid cells compared to ductal, acinar, beta, or alpha cells (tables S51 to S57).

### Monocle Analysis

Ductal, Alpha Major, Beta Major, Hybrid, Alpha/Beta, and Beta Minor cells were analyzed using the standard Monocle3 pipeline (*30, 56, 57*). Only genes within the pancreatic gene expression set (*22, 28*) were used for this analysis.

### CyTOF data collection, input files, and preprocessing

Flow CyTOF was performed as described previously (*22*). Briefly, after isolating the dissociated cells, barcoding was conducted for donors following the manufacturer’s protocol (Fluidigm, 101-0804 B1). Following barcoding, metal-conjugated antibody labeling was carried out in ‘FoxP3 permeabilization buffer’ (eBioscience, 00-8333) with 1% FBS (Hyclone, Cat# 7207) for 12 hours at 4°C at a concentration of up to 3 million cells per 300 μl of antibody cocktail, followed by twice washing with FoxP3 permeabilization buffer. Cells were then incubated with the DNA intercalator Iridium (Fluidigm, 201192A) at a dilution of 1:4,000 in 2% paraformaldehyde (Electron Microscopy Sciences, 15714) in DPBS (Corning, 21-031-CV) at RT for 1 hr. Mass cytometry data were acquired on CyTOF (Fluidigm). Flow CyTOF data analyses of endocrine cell composition was performed using the Cytobank implement (https://www.cytobank.org/).

Normalized FCS files were pre-processed prior to TooManyCells analysis and visualization using FlowJo Version 10.6.1 by gating all events on singlets according to event length and DNA content and then on live cells based on cisplatin exclusion. The Singlet/Live gated population was exported to a CSV file for TooManyCells analysis. Two dimensional plots were visualized for combinations of individual channels.

### TooManyCells clustering for CyTOF

TooManyCells was used to generate cell clades of CyTOF data. Cells with less than a total of 1e-16 signal were removed, leaving 6,945,575 cells. Upon inspection of protein levels across a tree with all cells (fig S17 a to c), endocrine and exocrine compartments were further subsetted leading to a refined analysis of 4,521,988 cells (Fig. 4A and 6A). Quantile normalization of the raw counts was used in the clustering step. The resulting tree was pruned by collapsing nodes with less than (7 MAD X median # cells in nodes) cells within them into their parent nodes.

### Imaging mass cytometry (IMC) analysis and Cell Segmentation

IMC was performed as described previously (*5*). Cell segmentation of all images was performed with the Vis software package (Visiopharm). All image channels were pre-processed with a 3×3 pixel median filter, then cells were segmented by applying a polynomial local linear parameter-based blob filter to the Iridium-193 DNA channel of each image to select objects representing individual nuclei. Identified nuclear objects were restricted to those greater than 10μm^2^, then dilated up to 7 pixels to approximate cell boundaries. Per-cell object mean pixel intensities were then exported for further analysis.

### TooManyCells clustering for IMC

TooManyCells was used to generate cell clades of IMC data. Cells with less than a total of 1e^-16^ signal were removed. Upon inspection of protein levels across a tree with all 1,170,001 cells (fig S17 f to h), endocrine and exocrine compartments were further subsetted, leading to the refined analysis of 130,428 cells for the investigation of the hybrid cell frequency (Fig. 4E). The full tree with 1,170,001 cells was used for the assessment of HLA-DR-expressing ductal cells (Fig 6F). Quantile normalization of the raw counts was used in the clustering step. The resulting tree was pruned by collapsing nodes with less than (5 MAD X median # cells in nodes) cells within them into their parent nodes. Subsetting of the tree was done with “--root-cut 3” to focus on node 3 in relevant analyses, with additional pruning of (3 MAD X median # cells in nodes).

### Cell-neighborhood analysis for IMC

Three labels were given to cells in the IMC neighborhood analysis: base, neighbor, and distant. Base cells originated from the chosen node, here node 16 in the node 3-focused IMC tree, or node 10 in the complete pruned tree which includes the former node Given the x- and y-coordinates from IMC per cell, each cell’s Euclidean distance to a base cell was calculated. If that distance was less than or equal to the chosen value, here 20 for the complete pruned tree, the cell was assigned the neighbor label. Otherwise the cell was designated as distant.

### IF and confocal microscopy

Tissues were fixed in 10% buffered formalin overnight, washed several times in PBS, then dehydrated through series of ethanol, embedded in paraffin and sectioned to 4-8um. Following deparaffinization through xylene and sequential rehydration, slides were subjected to heat antigen retrieval in a pressure cooker with Bulls Eye Decloaking buffer (Biocare). Slides were incubated in primary antibody overnight and secondary antibody conjugated to peroxidase and then developed using Tyramide Signal Amplification (TSA, Akoya Biosciences). Slides were counter stained with DAPI, mounted and imaged on Zeiss LSM800. Primary antibodies used for staining were Mouse anti-CK19 (Santa Cruz sc-6278) and Rabbit anti-HLA-DR (Abcam ab92511).

### Statistical analysis of box plots with Control, AAB+, and T1D donor states

The D’Agostino & Pearson omnibus normality test was used to assess whether the data from each group was normally distributed. If any group failed the D’Agostino & Pearson omnibus normality test, the Kruskal-Wallis test was applied. If none of the groups failed the D’Agostino & Pearson omnibus normality test, the one-way ANOVA test was applied.

### Statistical analysis of cellular neighborhoods

Differential marker expression significance for neighbors in the IMC analysis was determined using permutation tests. For each marker, the distribution of that marker value for each of the designated n neighbors was compared against 100 distributions derived from n random cells across the entire IMC tree. The resulting p-value was calculated by the ratio of the number of permutations that had a lower median marker value than the observed marker value to the total number of permutations. If this value was < 0.5, the value was subtracted from 1 to switch directionality (number of permutations with a higher median value). To account for the two-tailed test, this value was multiplied by 2 for the final p-value calculation.

---

**Figure S1:**
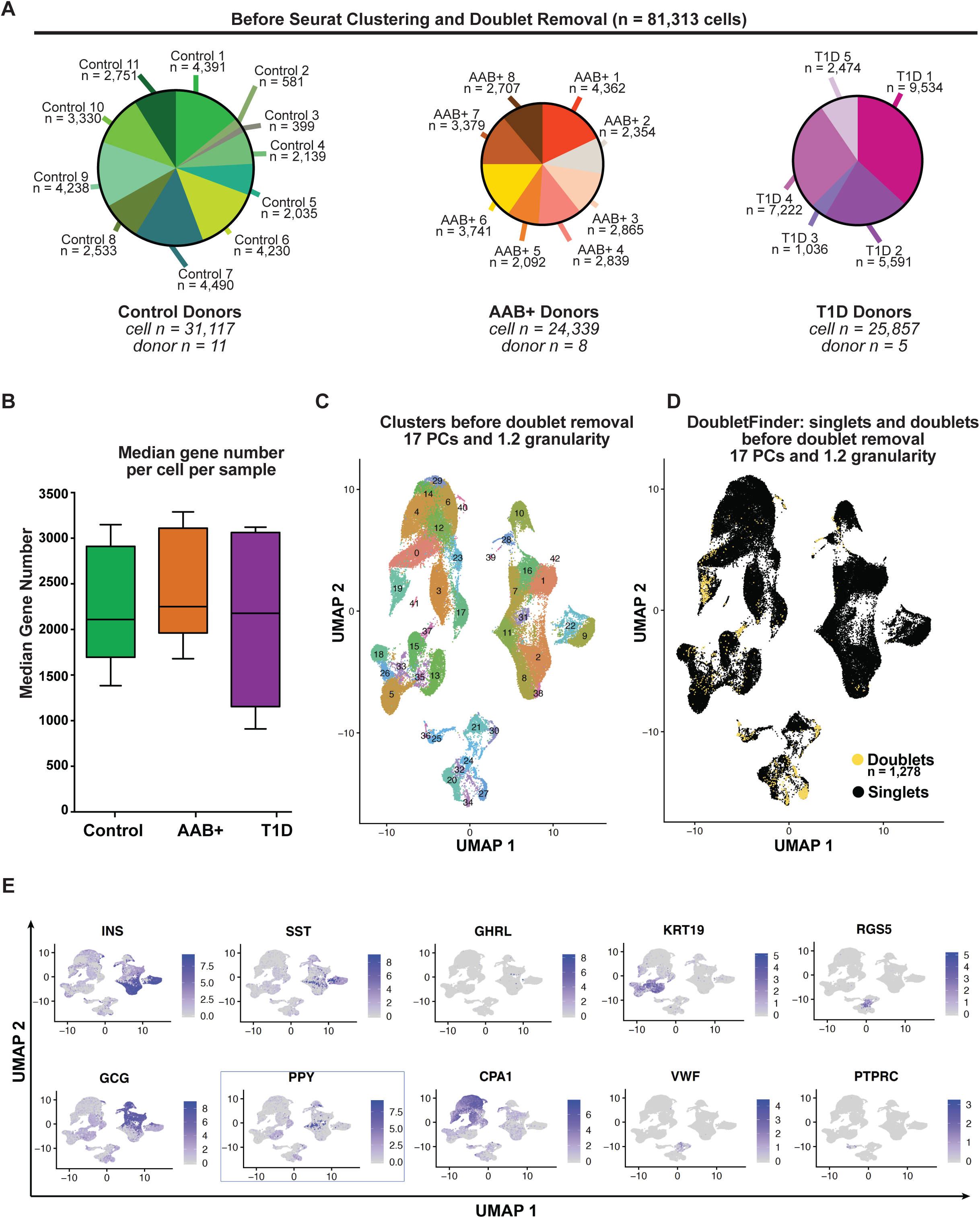
Cell numbers and clustering before complete filtering. (A) Pie chart displaying the cell numbers/proportions of each individual donor per donor type. (B) Box plot displaying the average gene number per cell per donor type. (C) UMAP visualization of cell clusters for all cells. (D) Doublets and singlets, as identified using DoubletFinder, across cell clusters visualized by UMAP. (E) UMAP visualization of the normalized gene expression counts of each canonical gene marker of each major cell type.

**Figure S2:**
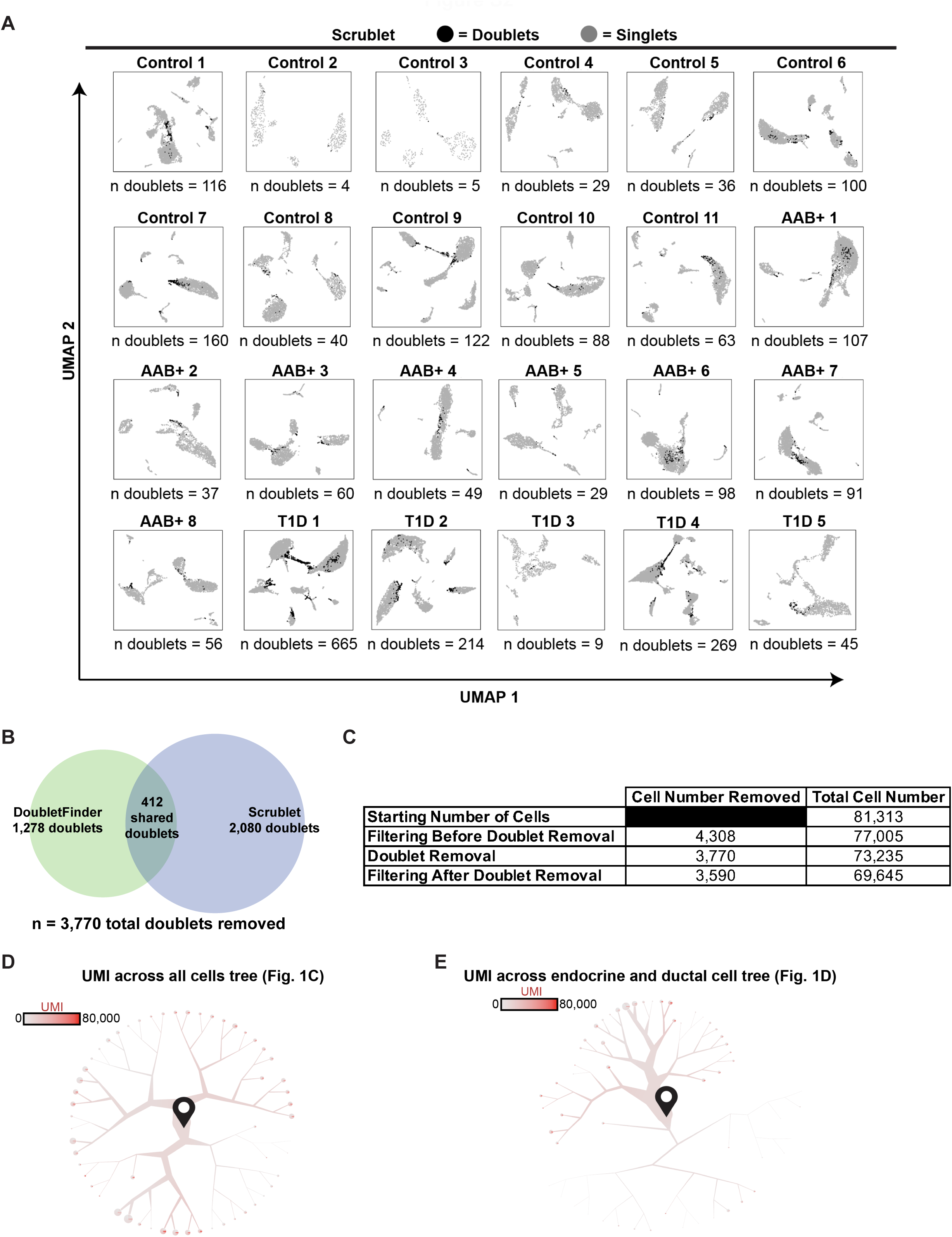
Doublet removal and UMI counts. (A) Doublets and singlets, as identified using Scrublet, across cell clusters visualized by UMAP per individual. (B) Venn diagram indicating the number of cells deemed doublets by DoubletFinder and Scrublet, as well as cells that were commonly identified by both approaches. (C) Table indicating the number of cells removed and the resulting total cell number for each step of filtering. (D) Unique molecular identifier (UMI) counts per cell projected across the dendrogram visualization and clustering of all cells from Figure 1C. Pie charts at the end of the branches display the breakdown of UMI counts per cell within that terminal cluster. Cells begin at the start pin symbol, and from there are partitioned based on similarities and differences in gene expression. (E) UMI counts per cell projected across the dendrogram visualization and clustering of ductal and endocrine cells from Figure 1D. Pie charts at the end of the branches display the breakdown of UMI counts per cell within that terminal cluster. Cells begin at the start pin symbol, and from there are partitioned based on similarities and differences in gene expression.

**Figure S3:**
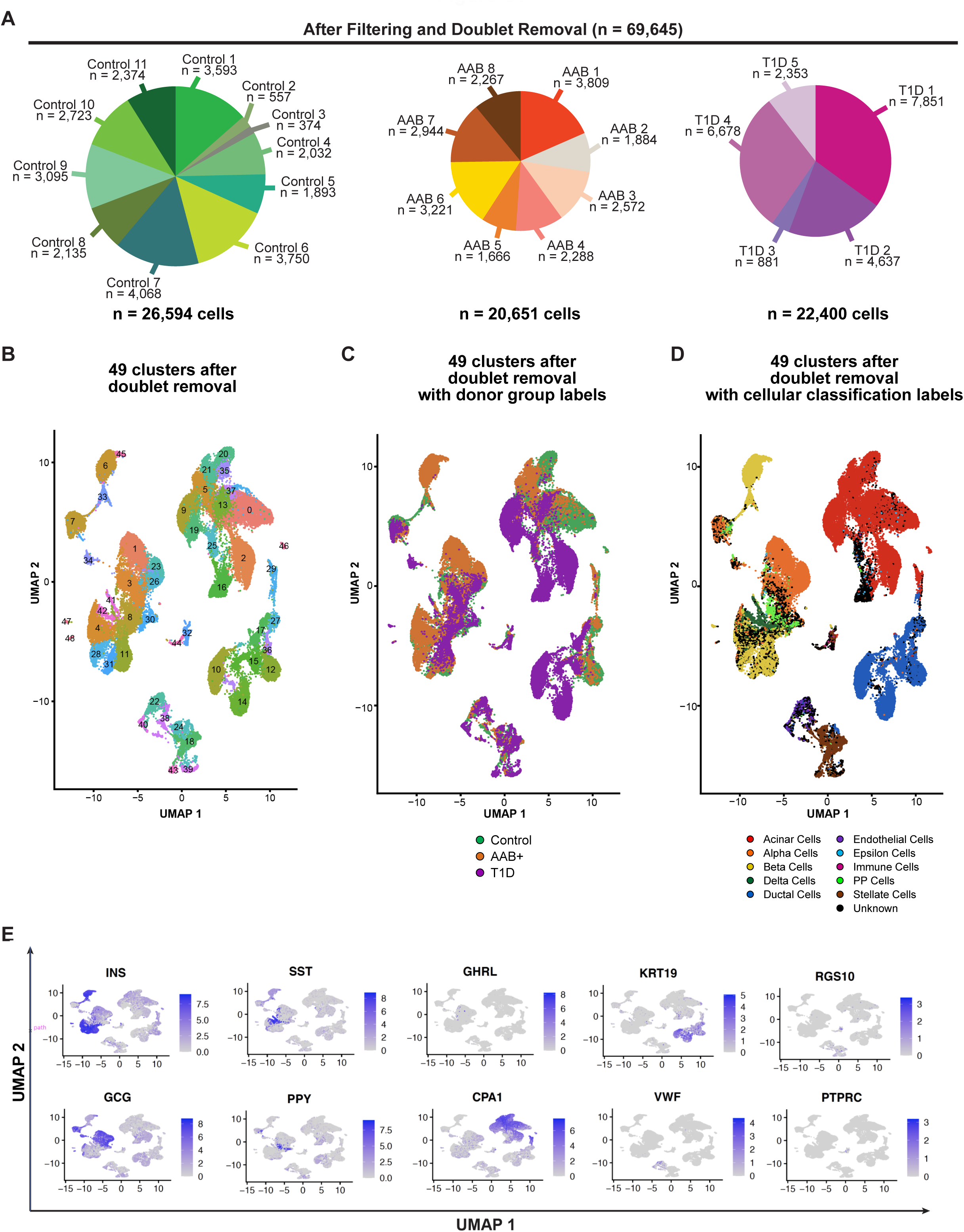
Cell numbers and clustering after complete filtering. (A) Pie chart displaying the cell numbers/proportions of each individual donor per donor type. (B) UMAP visualization of cell clusters for all cells. (C) UMAP visualization donor groups across clusters for all cells. (D) UMAP visualization of Garnett cellular classifications across clusters for all cells. (E) UMAP visualization of the normalized gene expression counts of each canonical gene marker of each major cell type.

**Figure S4:**
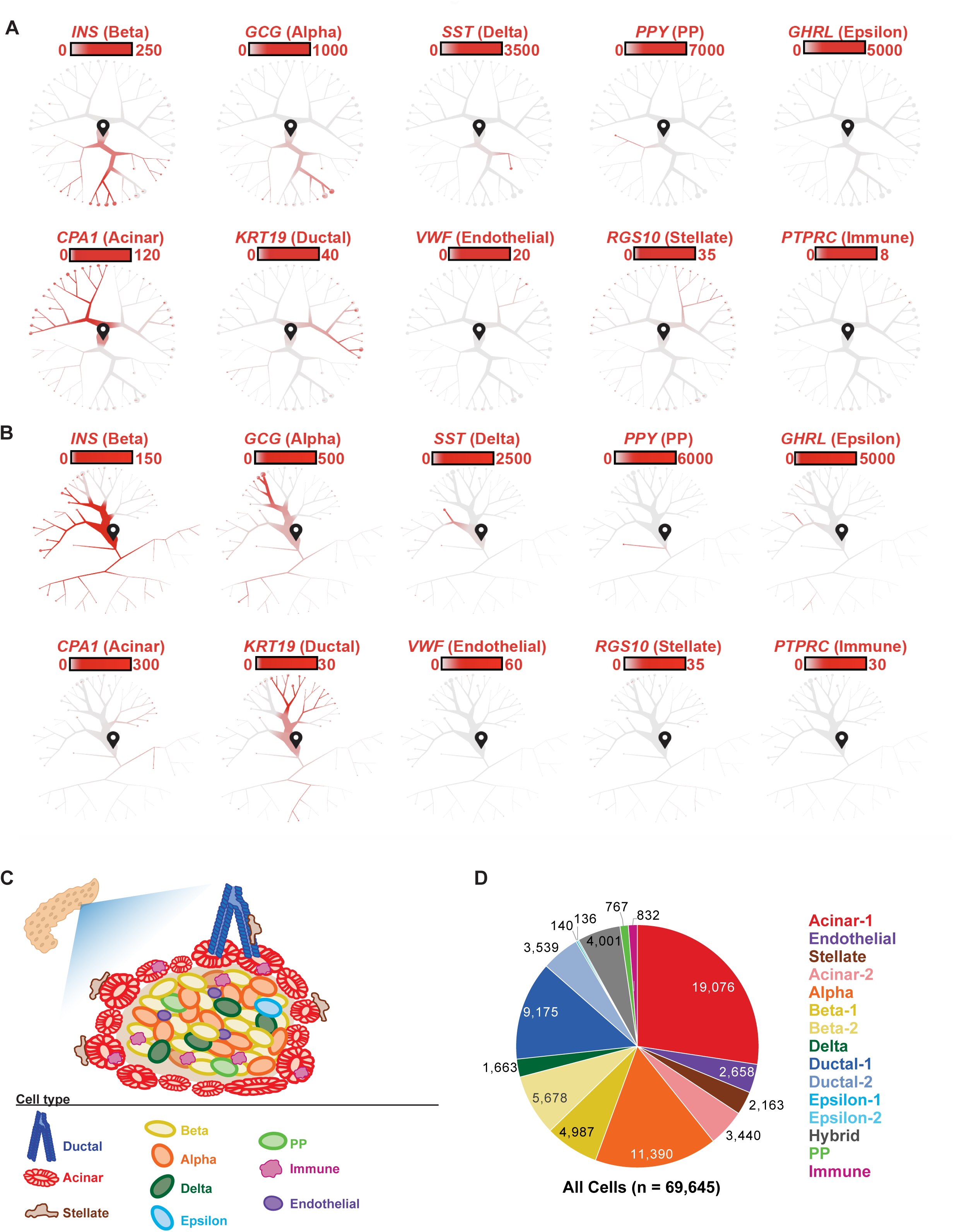
Marker gene expression confirms canonical cell types. (A) Dendrograms highlighting the expression of each canonical gene marker of each major cell type across the dendrogram of all cells in Figure 1C. Scale bars represent normalized transcript numbers. Cells begin at the start pin symbol, and from there are partitioned based on similarities and differences in gene expression. (B) Dendrograms highlighting the expression of each canonical gene marker of each major cell type across the dendrogram of ductal and endocrine cells in Figure 1D. Scale bars represent normalized transcript numbers. Cells begin at the start pin symbol, and from there are partitioned based on similarities and differences in gene expression. (C) Schematic of the human pancreatic islet anatomy and major cell types. (D) Pie chart displaying the cell numbers/proportions of each cell type defined in Figure 1, C and D.

**Figure S5:**
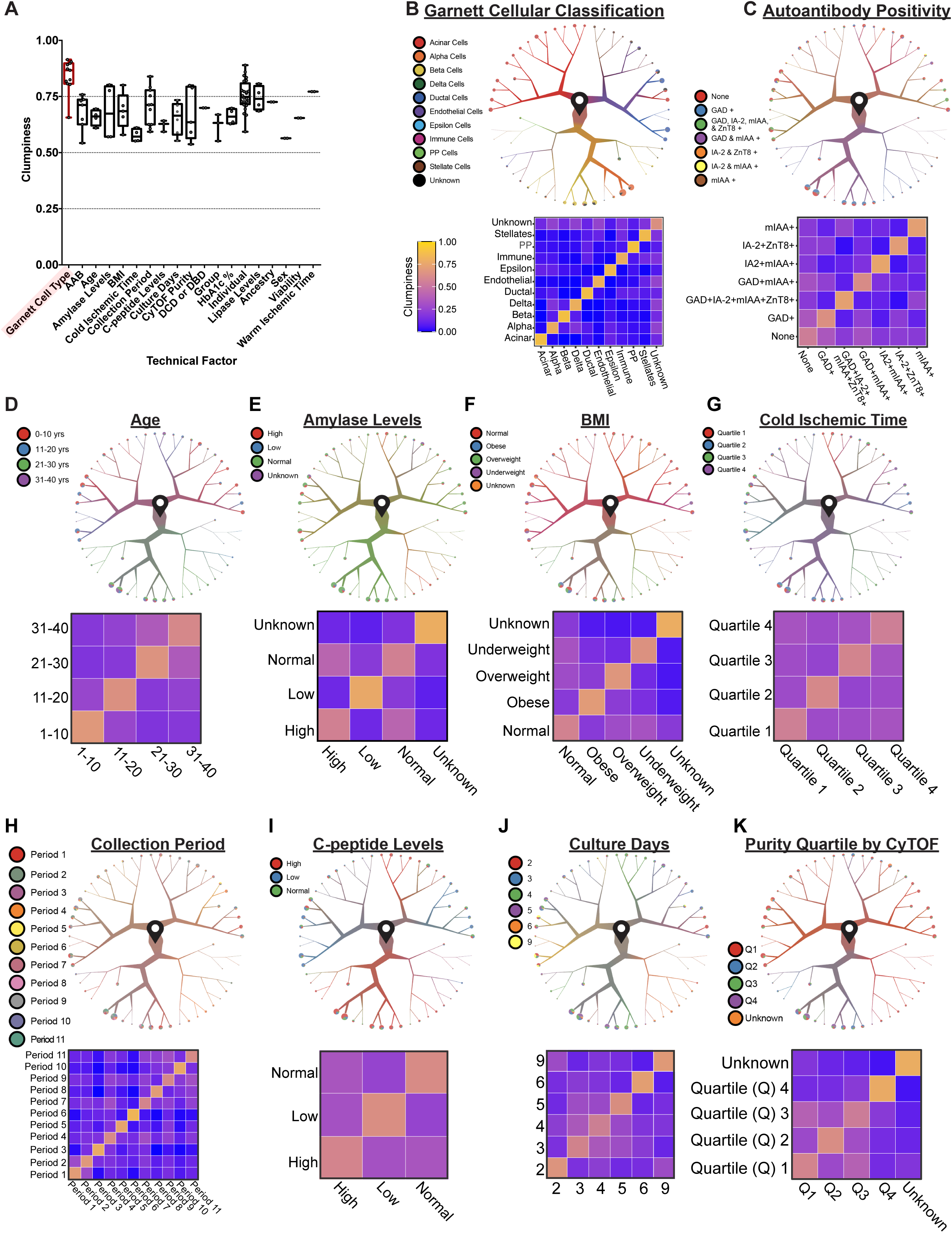
Technical factors or biological factors do not drive cell segregation. (A) Box plot displaying clumpiness measure across biological and technical factors. (B-K) Cells begin at the start pin symbol, and from there are partitioned based on similarities and differences in gene expression. (Top) Technical or biological factors projected across the dendrogram visualization and clustering of all cells from Figure 1C. Pie charts at the end of the branches display the breakdown of technical or biological factors within that terminal cluster. (Bottom) Quantification of the relationship between different technical or biological factors using the clumpiness measure as determined from the dendrograms. Clumpiness is a measure of aggregation of labels within a dendrogram. Here, each leaf of the dendrogram contains a collection of labels (technical or biological covariates). The more labels group together within the dendrogram, the higher the clumpiness value. When using hierarchy in biological systems to elucidate relationships between metadata, the distribution of metadata labels within the hierarchy may exhibit different levels of aggregation—uniform, random, or clumped. Our strategy for estimating the impact of technical and biological variance on substructures beyond cell type identity was to use a measure called “clumpiness”. Clumpiness is a measure for finding the level of aggregation between labels distributed among the leaves of a hierarchical tree and extensively measures the relationships between metadata. We have shown that our measure is robust to the cell number, data size, and label size, and thus is scale invariant. In addition, our measure is generalizable to more than two labels and is efficiently computable and maintainable. In the revised manuscript Supplemental Figures S5 and S6, we demonstrate that although cells from individual donors tend to cluster with each other as expected, no particular feature affected cell type identity definition in our analytical approach.

**Figure S6:**
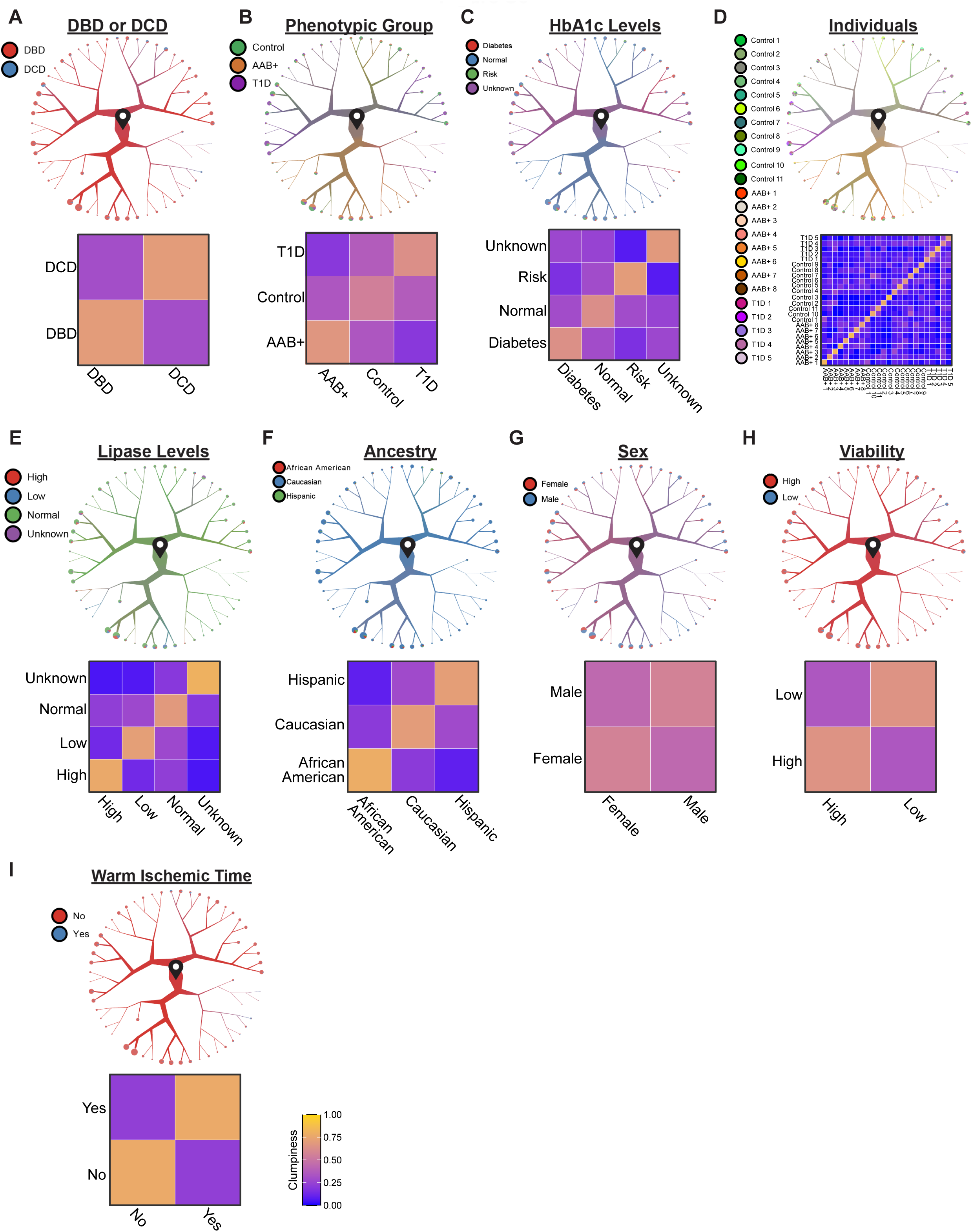
Technical factors or biological factors do not drive cell segregation. (A-I) Cells begin at the start pin symbol, and from there are partitioned based on similarities and differences in gene expression. (Top) Technical or biological factors projected across the dendrogram visualization and clustering of all cells from Figure 1C. Pie charts at the end of the branches display the breakdown of technical or biological factors within that terminal cluster. (Bottom) Quantification of the relationship between different technical or biological factors using the clumpiness measure as determined from the dendrograms. Clumpiness is a measure of aggregation of labels within a dendrogram. Here, each leaf of the dendrogram contains a collection of labels (technical or biological covariates). The more labels group together within the dendrogram, the higher the clumpiness value. When using hierarchy in biological systems to elucidate relationships between metadata, the distribution of metadata labels within the hierarchy may exhibit different levels of aggregation—uniform, random, or clumped. Our strategy for estimating the impact of technical and biological variance on substructures beyond cell type identity was to use a measure called “clumpiness”. Clumpiness is a measure for finding the level of aggregation between labels distributed among the leaves of a hierarchical tree and extensively measures the relationships between metadata. We have shown that our measure is robust to the cell number, data size, and label size, and thus is scale invariant. In addition, our measure is generalizable to more than two labels and is efficiently computable and maintainable. In the revised manuscript Supplemental Figures S5 and S6, we demonstrate that although cells from individual donors tend to cluster with each other as expected, no particular feature affected cell type identity definition in our analytical approach.

**Figure S7:**
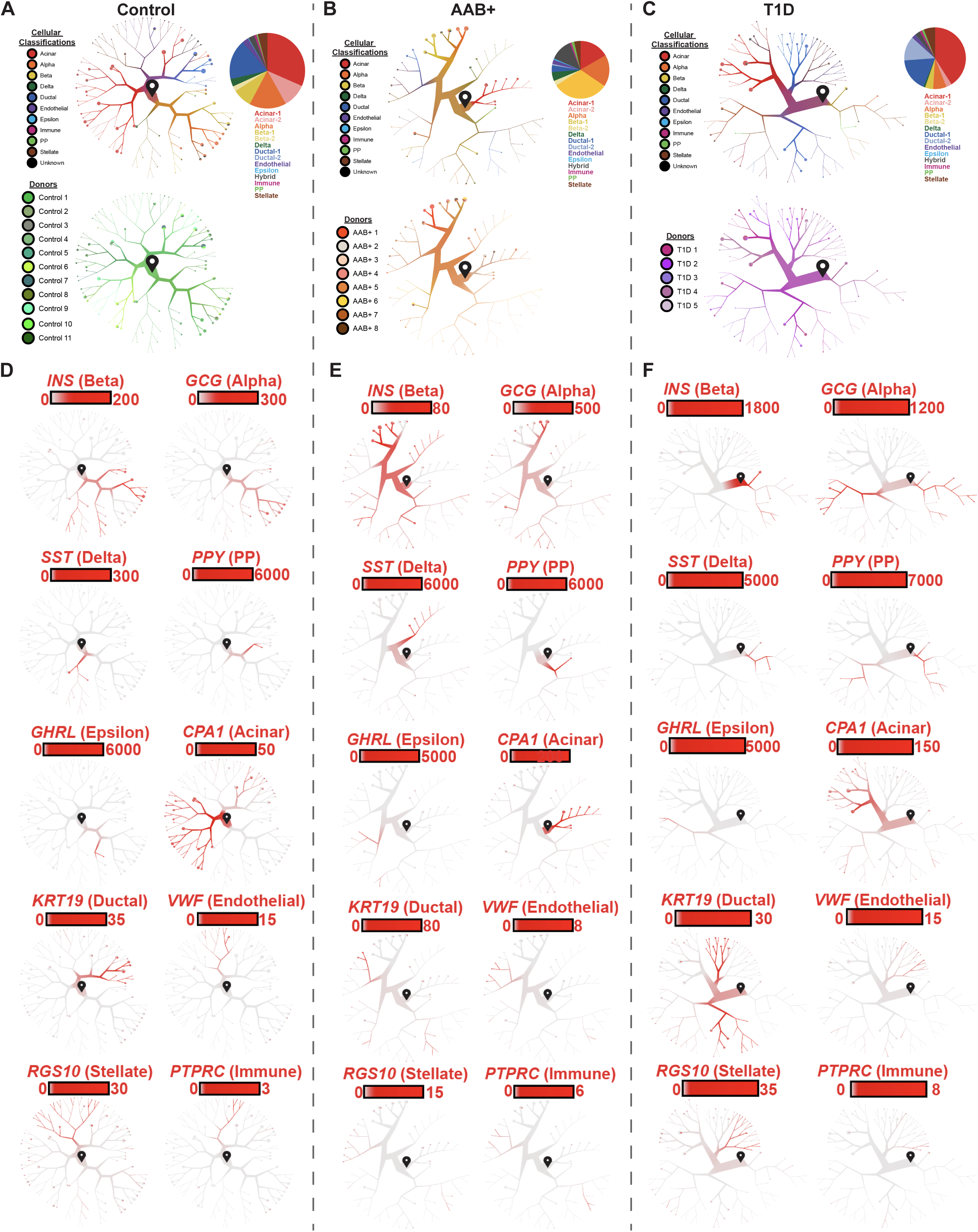
Discernment of cell types when clustering cells from each disease state independently. (A-C) Cells begin at the start pin symbol, and from there are partitioned based on similarities and differences in gene expression. Dendrogram visualization and clustering of all cells from each of the donor states independently. Cells begin at the start pin symbol. Garnett cellular classifications (Top Left) and individual donor (Bottom Left) labels across the dendrogram. (Right) Pie chart displaying the cell proportions of each cell type. (D-F) Dendrogram visualization of the expression of each canonical gene marker of each major cell type across the dendrograms from A-C. Scale bars represent normalized transcript numbers.

**Figure S8:**
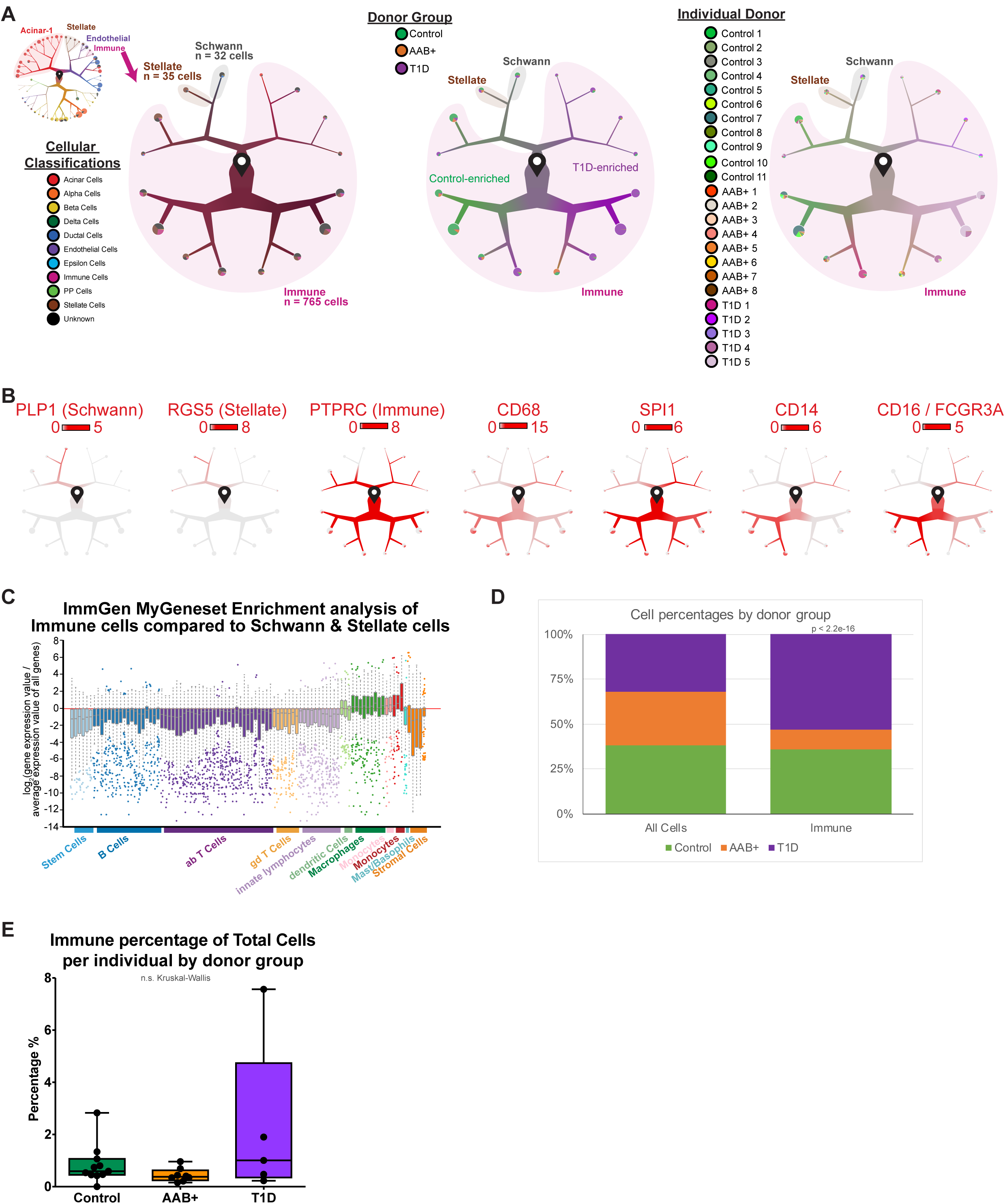
Discernment of cell types when clustering cells from each disease state independently. (A) Cells begin at the start pin symbol, and from there are partitioned based on similarities and differences in gene expression. (Inset) Dendrogram visualization and clustering of all cells from Figure 1C. Highlighted in pink is the immune cell cluster, which was subsetted and re-clustered here. (Main) Dendrogram visualization of clustered immune cells (along with small clusters of Stellate and Schwann cells) with Garnett cellular classification (left), donor group (middle), or individual donor (right) designations across the tree. (B) Dendrograms highlighting the expression of each canonical gene marker of each relevant cell type across the dendrogram of immune cells in Figure S8A. Scale bars represent normalized transcript numbers. (C) Gene set enrichment of ImmGen cell types for the genes upregulated in the *PTPRC* high immune cell cluster (pink in fig. S8A) when compared to the closely transcriptionally related stellate (brown in fig. S8A) plus Schwann cells (grey in fig. S8A). (D) Bar graph displaying the proportion of immune cells from each donor group. P-value presented is the result of the Chi-squared test. (E) Box plots displaying immune cell percentage of total cells per individual across donor groups.

**Figure S9:**
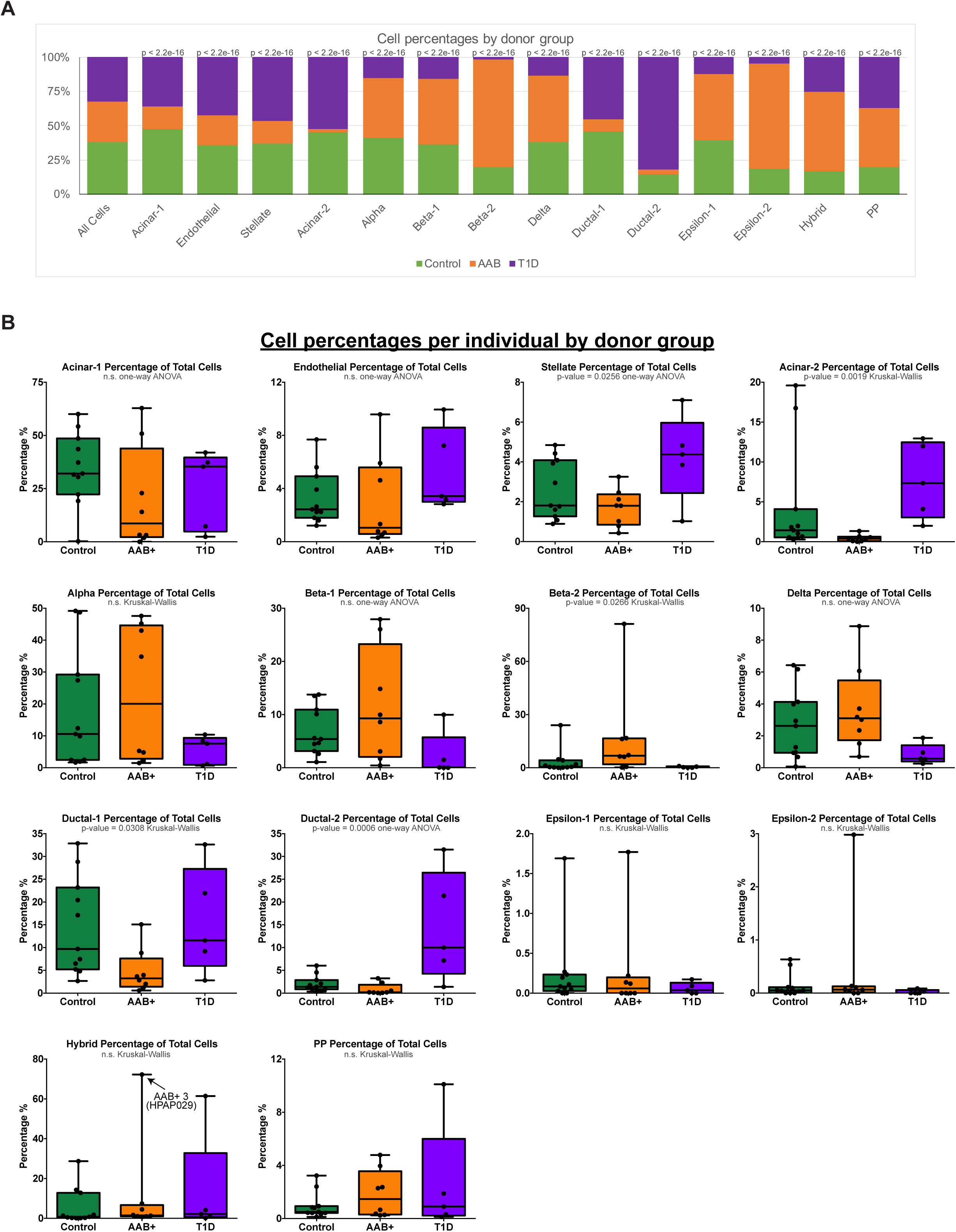
Discernment of cell types when clustering cells from each disease state independently. (A) Bar graph displaying the proportion of cells from each donor group for all major pancreatic cell types. P-values presented are the result of the Chi-squared test. (B) Box plots displaying cell type percentage of total cells per individual across donor groups for all major pancreatic cell types.

**Figure S10:**
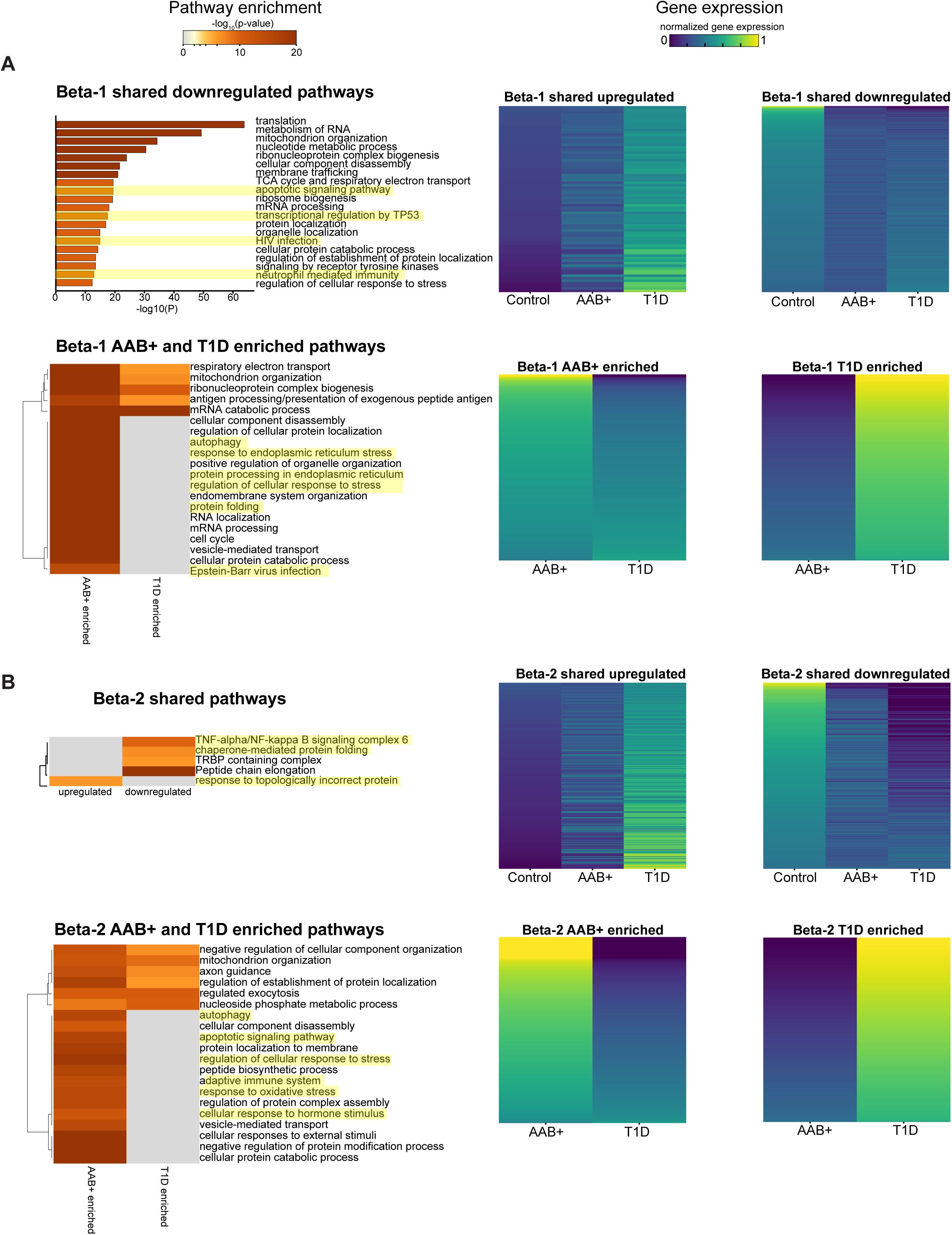
Gene and gene ontology pathways that are shared and different across disease states in beta cells. (A-B) (Left) For each cell type, displayed are gene ontology pathways that are shared across T1D and AAB+ cells when compared to Control cells (top) or pathways that are differently enriched in T1D cells vs AAB+ cells (bottom). The top 20 clusters are displayed and a stringent cut-off of p ≤ 1e-6 was applied to determine significant gene ontology pathways. Pathways highlighted in yellow are displayed in Figure 2. (Right) Heatmaps displaying the degree of gene expression changes of genes (rows) that are shared (top) or differential (bottom) across AAB+ and T1D disease states.

**Figure S11:**
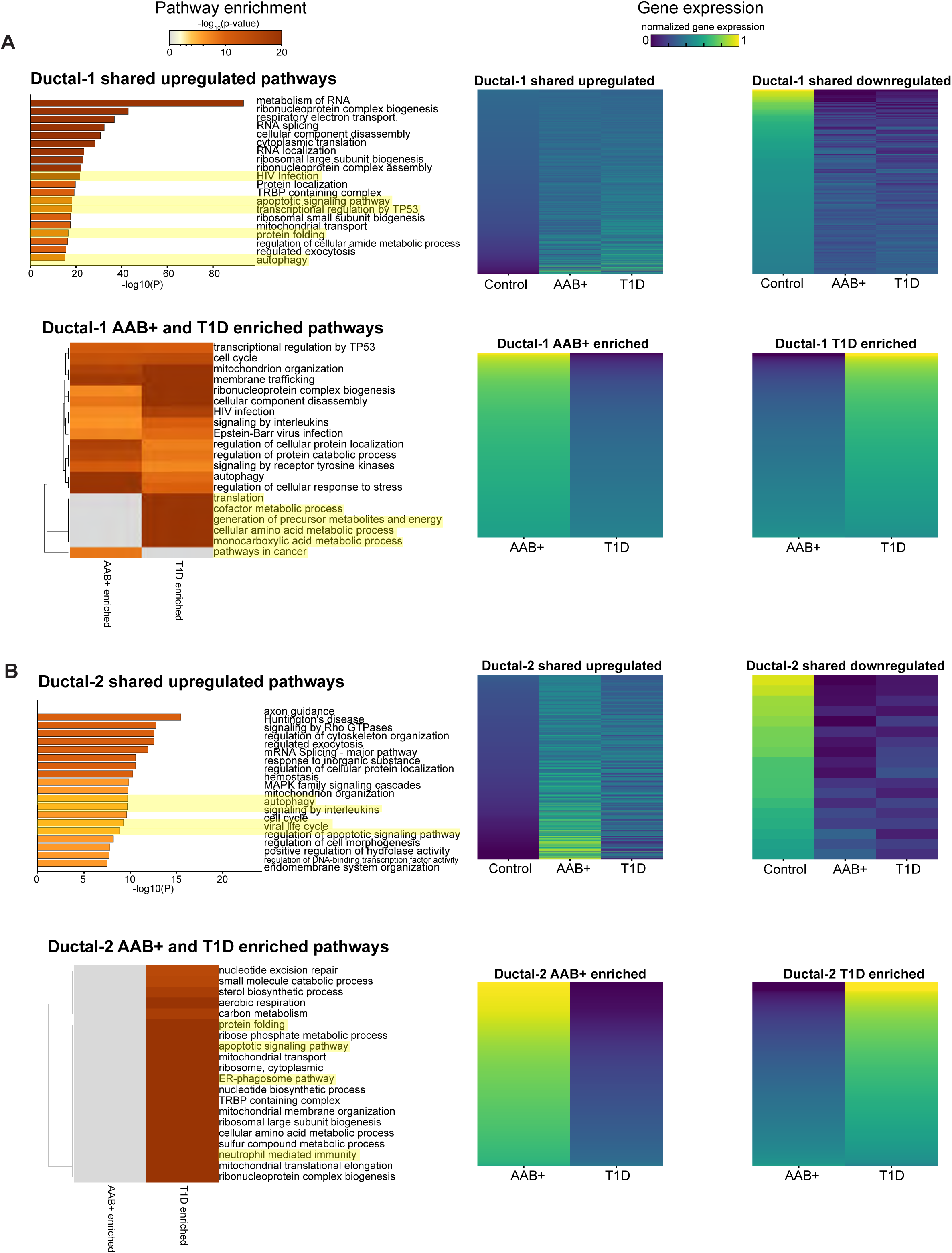
Gene and gene ontology pathways that are shared and different across disease states in ductal cells. (A-B) (Left) For each cell type, displayed are gene ontology pathways that are shared across T1D and AAB+ cells when compared to Control cells (top) or pathways that are differently enriched in T1D cells vs AAB+ cells (bottom). The top 20 clusters are displayed and a stringent cut-off of p ≤ 1e-6 was applied to determine significant gene ontology pathways. Pathways highlighted in yellow are displayed in Figure 2. (Right) Heatmaps displaying the degree of gene expression changes of genes (rows) that are shared (top) or differential (bottom) across AAB+ and T1D disease states.

**Figure S12:**
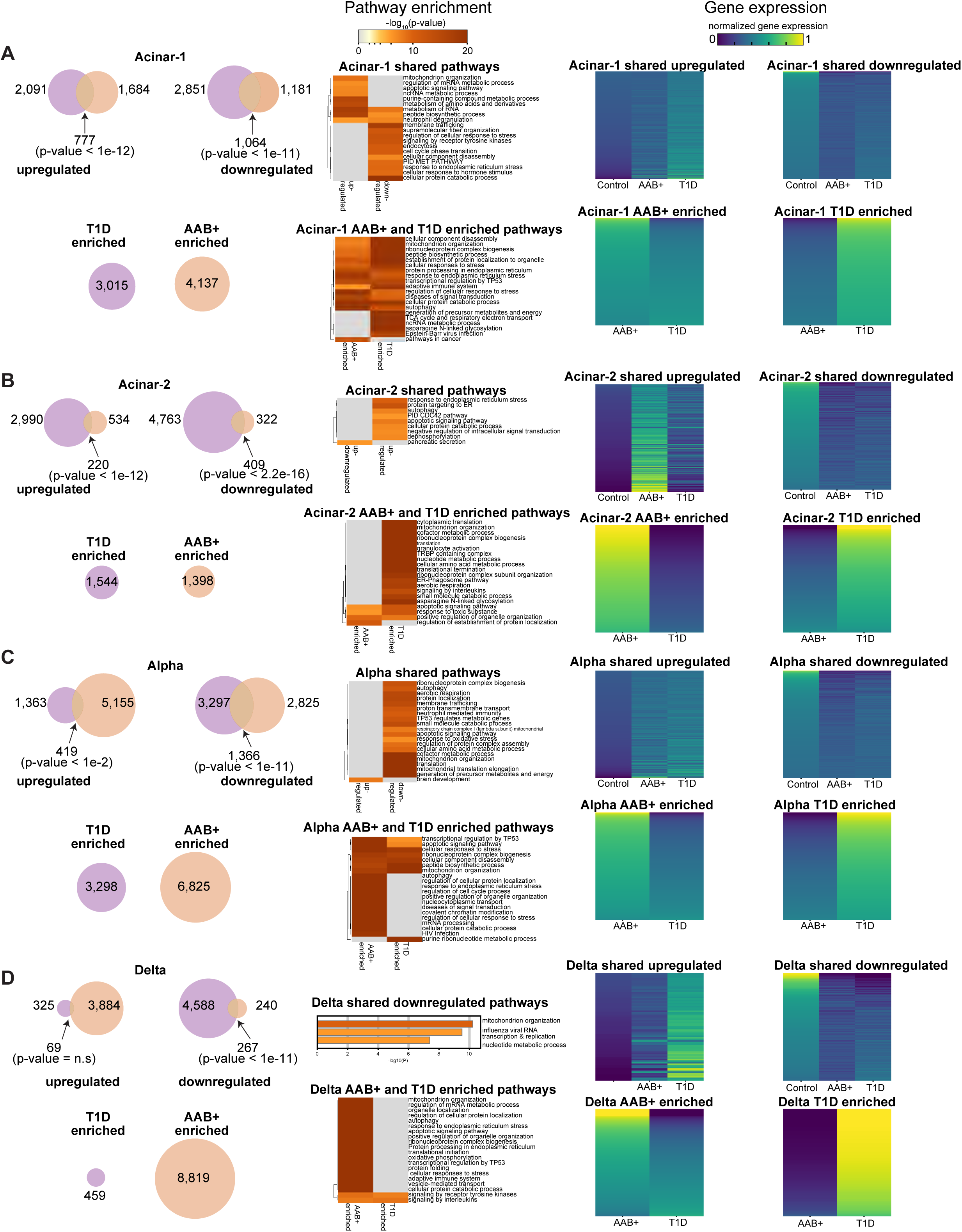
Gene and gene ontology pathways that are shared and different across disease states in Acinar-1, Acinar-2, Alpha, and Delta cells. (A-D) (Left) For each cell type, Venn diagrams indicate the numbers of upregulated and downregulated genes, as well as overlapping genes, across the two donor states. Circles indicate the numbers of genes that are ‘T1D enriched’ or ‘AAB enriched’. P-values presented are the results of hypergeometric CDF tests (one-tailed test for overrepresentation). (Middle) For each cell type, displayed are gene ontology pathways that are shared across T1D and AAB+ cells when compared to Control cells (top) or pathways that are differently enriched in T1D cells vs AAB+ cells (bottom). The top 20 clusters are displayed and a stringent cut-off of p ≤ 1e-6 was applied to determine significant gene ontology pathways. (Right) Heatmaps displaying the degree of gene expression changes of genes (rows) that are shared (top) or differential (bottom) across AAB+ and T1D disease states.

**Figure S13:**
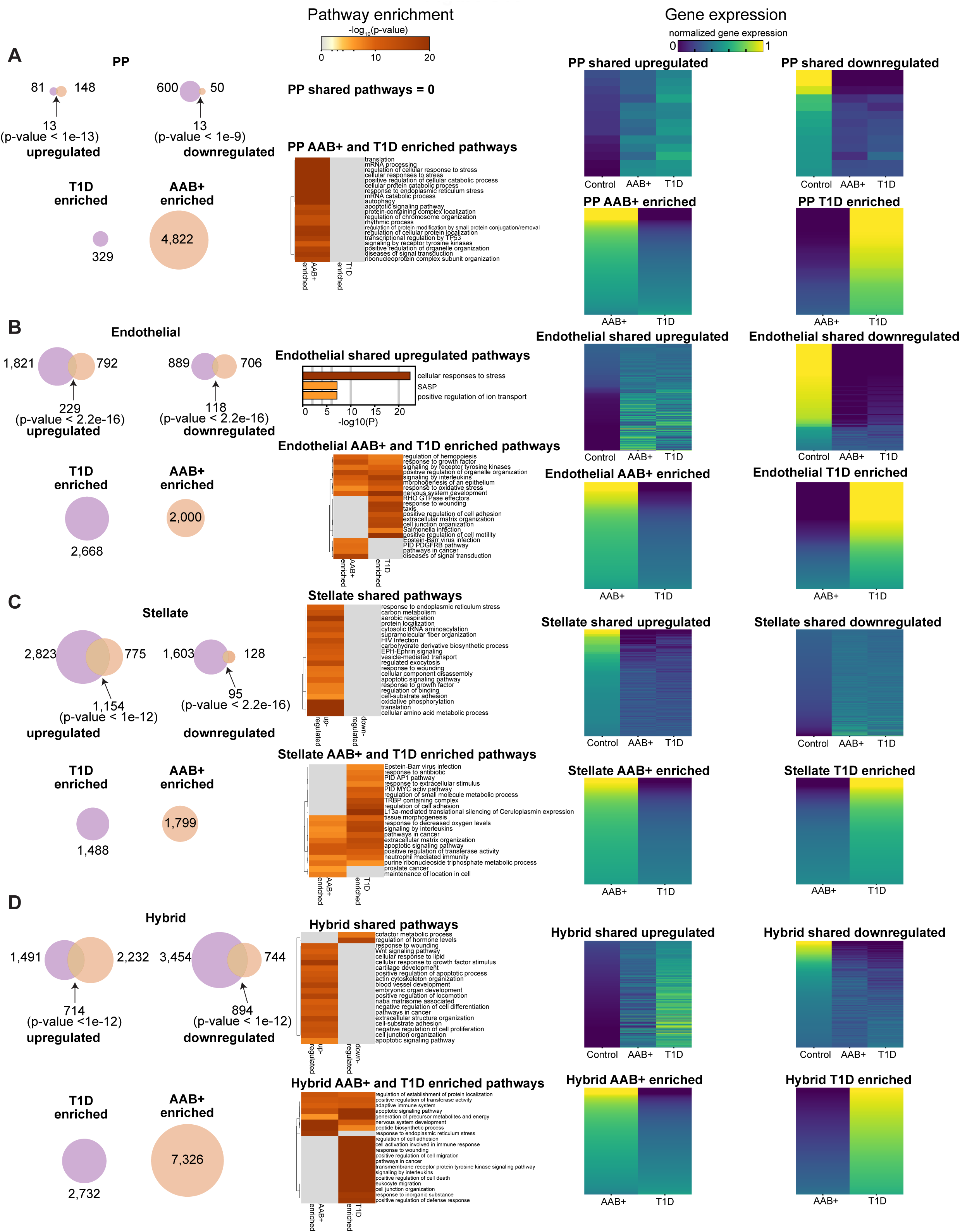
Gene and gene ontology pathways that are shared and different across disease states in PP, Endothelial, Stellate, and Hybrid cells. (A-D) (Left) For each cell type, Venn diagrams indicate the numbers of upregulated and downregulated genes, as well as overlapping genes, across the two disease states. Circles indicate the numbers of genes that are ‘T1D enriched’ or ‘AAB enriched’. P-values presented are the results of hypergeometric CDF tests (one-tailed test for overrepresentation). (Middle) For each cell type, displayed are gene ontology pathways that are shared across T1D and AAB+ cells when compared to Control cells (top) or pathways that are differently enriched in T1D cells vs AAB+ cells (bottom). The top 20 clusters are displayed and a stringent cut-off of p ≤ 1e-6 was applied to determine significant gene ontology pathways. (Right) Heatmaps displaying the degree of gene expression changes of genes (rows) that are shared (top) or differential (bottom) across AAB+ and T1D disease states.

**Figure S14:**
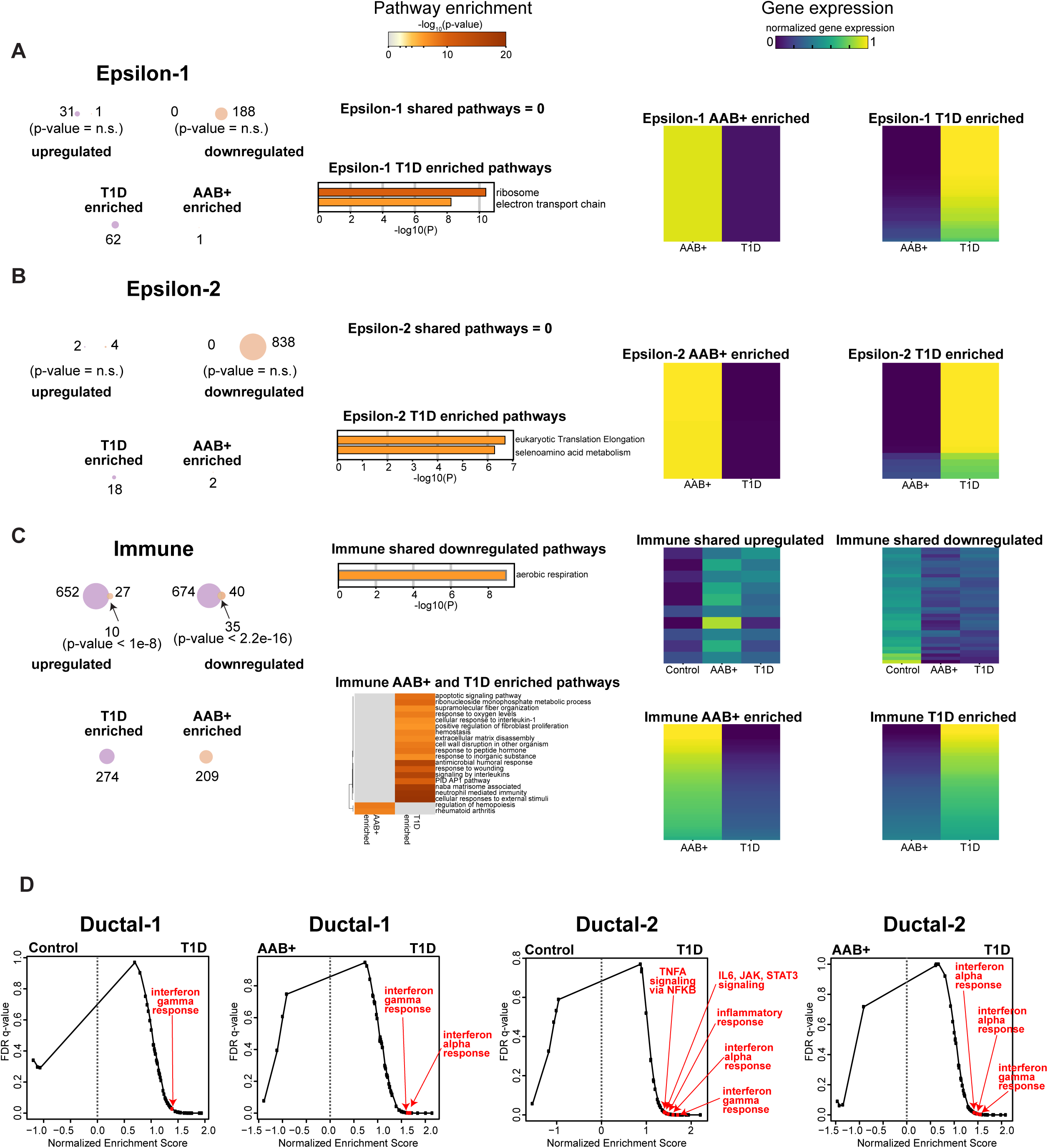
Gene and gene ontology pathways that are shared and different across disease states in Epsilon-1, Epsilon-2, and Immune cells. (A-C) (Left) For each cell type, Venn diagrams indicate the numbers of upregulated and downregulated genes, as well as overlapping genes, across the two disease states. Circles indicate the numbers of genes that are ‘T1D enriched’ or ‘AAB enriched’. P-values presented are the results of hypergeometric CDF tests (one-tailed test for overrepresentation). (Middle) For each cell type, displayed are gene ontology pathways that are shared across T1D and AAB+ cells when compared to Control cells (top) or pathways that are differently enriched in T1D cells vs AAB+ cells (bottom). The top 20 clusters are displayed and a stringent cut-off of 1e-6 was applied to determine significant gene ontology pathways. (Right) Heatmaps displaying the degree of gene expression changes of genes (rows) that are shared (top) or differential (bottom) across AAB+ and T1D disease states. (D) GSEA analysis plots of FDR q-value vs Normalized Enrichment Score. For both ductal populations, Ductal-1 and Ductal-2, T1D cells were compared to AAB+ or Control cells to determine differentially enriched gene sets. Demarcated in red and labeled are signatures of interest.

**Figure S15:**
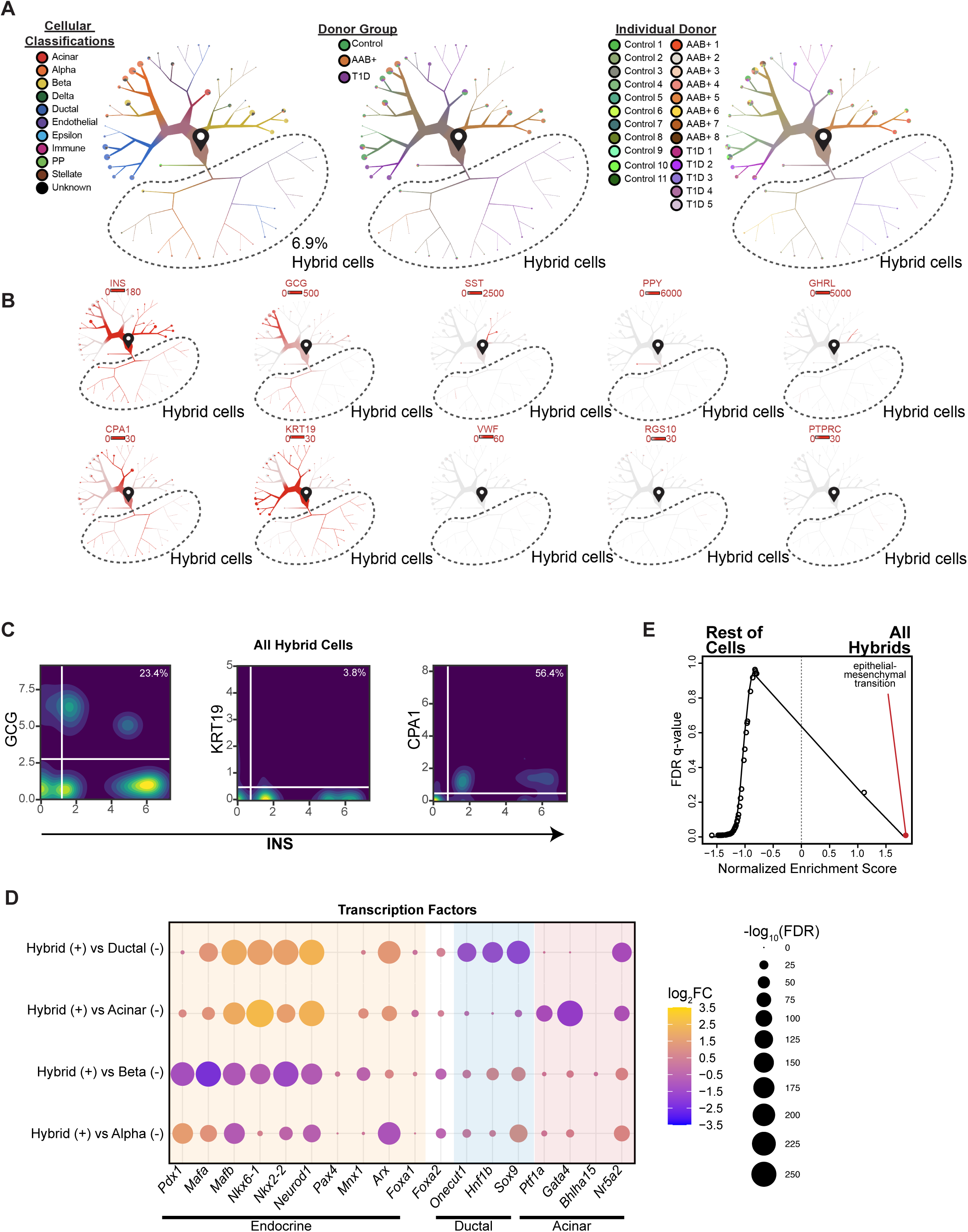
The hybrid cellular state exists across multiple donors and has unique transcriptional features. (A) Cells begin at the start pin symbol, and from there are partitioned based on similarities and differences in gene expression. Dendrogram visualization and clustering of ductal and endocrine cells, without the dominant donor that contributes to Hybrid cells, AAB+ donor #3. Cells begin at the start pin symbol. Pie charts at the end of the branches display the breakdown of Garnett cellular classification (left), the donor group (middle), or individual donors (right) within that terminal cluster. Circled in black is the Hybrid cluster. (B) Dendrograms highlight the expression of each canonical gene marker of each major cell type across the dendrogram in fig. S15A. Scale bars represent normalized transcript numbers. (C) Co-expression analysis of *INS* and the relevant gene indicative of each subpopulation of Hybrid cells on an individual cell level across all Hybrid cells (Ductal-like Hybrids plus Endocrine-like Hybrids plus Acinar-like Hybrids). For each gene, values represent normalized transcript numbers. Percentages in the upper right-hand corner indicate the percentage of co-expressing cells. (D) Dot plot displaying the log_2_(Fold Change (FC)) (circle color) and -log_10_(FDR) (circle size) of transcription factors (TFs) that are important for endocrine, ductal, and acinar cell fates across four comparisons: Hybrid versus Ductal (Hybrid/(Ductal-1 plus Ductal-2)), Hybrid versus Acinar (Hybrid/Acinar-2), Hybrid versus Beta (Hybrid/(Beta-1 plus Beta-2)), and Hybrid versus Alpha (Hybrid/Alpha). *FOXA2* is a transcription factor important in both endocrine and ductal cell fates. (E) GSEA analysis plots of FDR q-value vs Normalized Enrichment Score. All Hybrid cells (Ductal-like Hybrids plus Endocrine-like Hybrids plus Acinar-like Hybrids) were compared to all other cell populations combined in the ductal and endocrine dendrogram to determine differentially enriched gene sets. Demarcated in red and labeled is a signature of interest.

**Figure S16:**
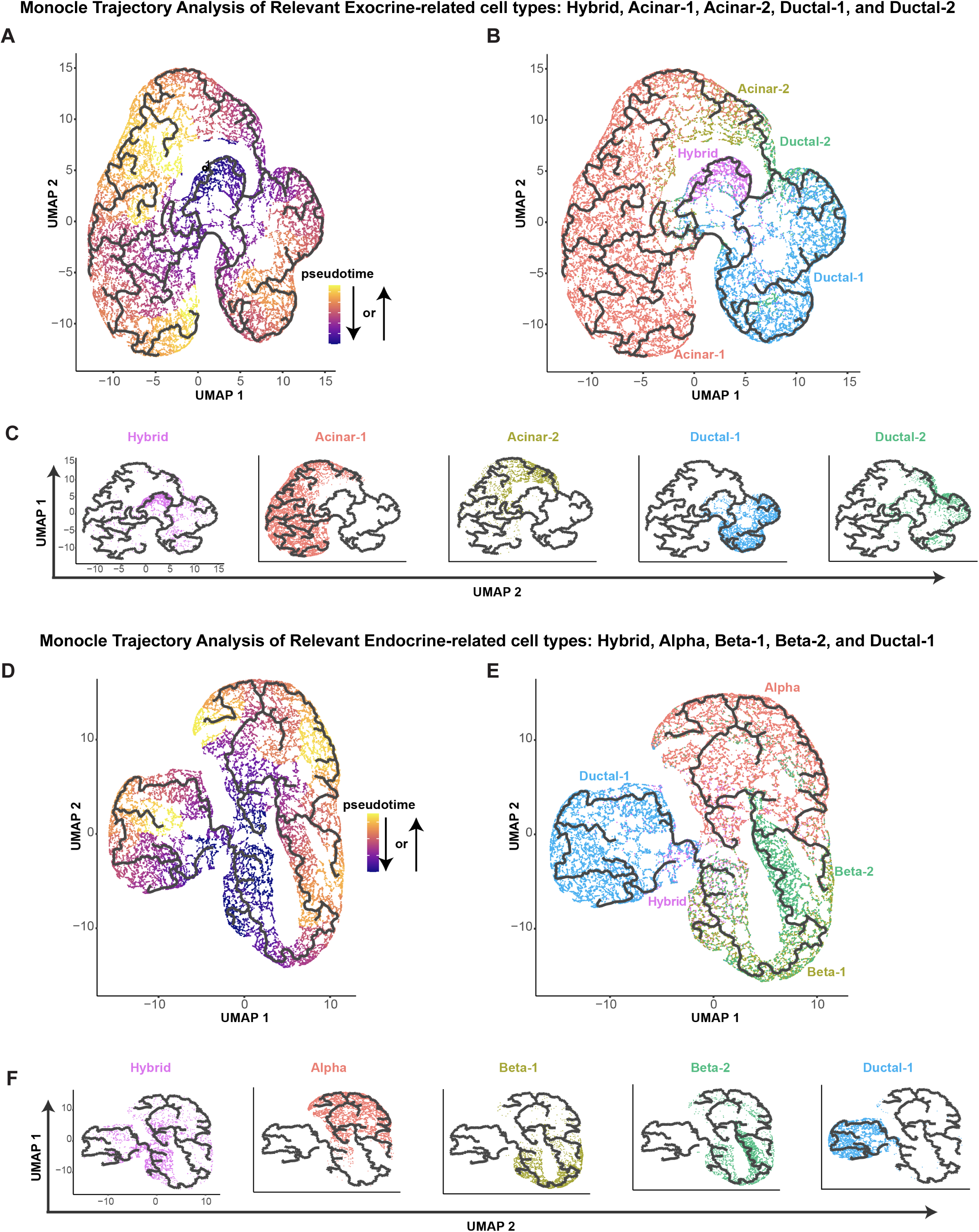
Monocle trajectory analysis. (A-C) Monocle trajectory analysis of exocrine-related cell types: Hybrid, Acinar-1, Acinar-2, Ductal-1, and Ductal-2. Dark grey line indicates a possible cellular trajectory. Cells are labeled by their pseudotime (A), all cellular types together (B), or individual cellular types separately (C). (D-E) Monocle trajectory analysis of endocrine-related cell types: Hybrid, Alpha, Beta-1, Beta-2, and Ductal-1. Dark grey line indicates a possible cellular trajectory. Cells are labeled by their pseudotime (A), all cellular types together (B), or individual cellular types separately (C).

**Figure S17:**
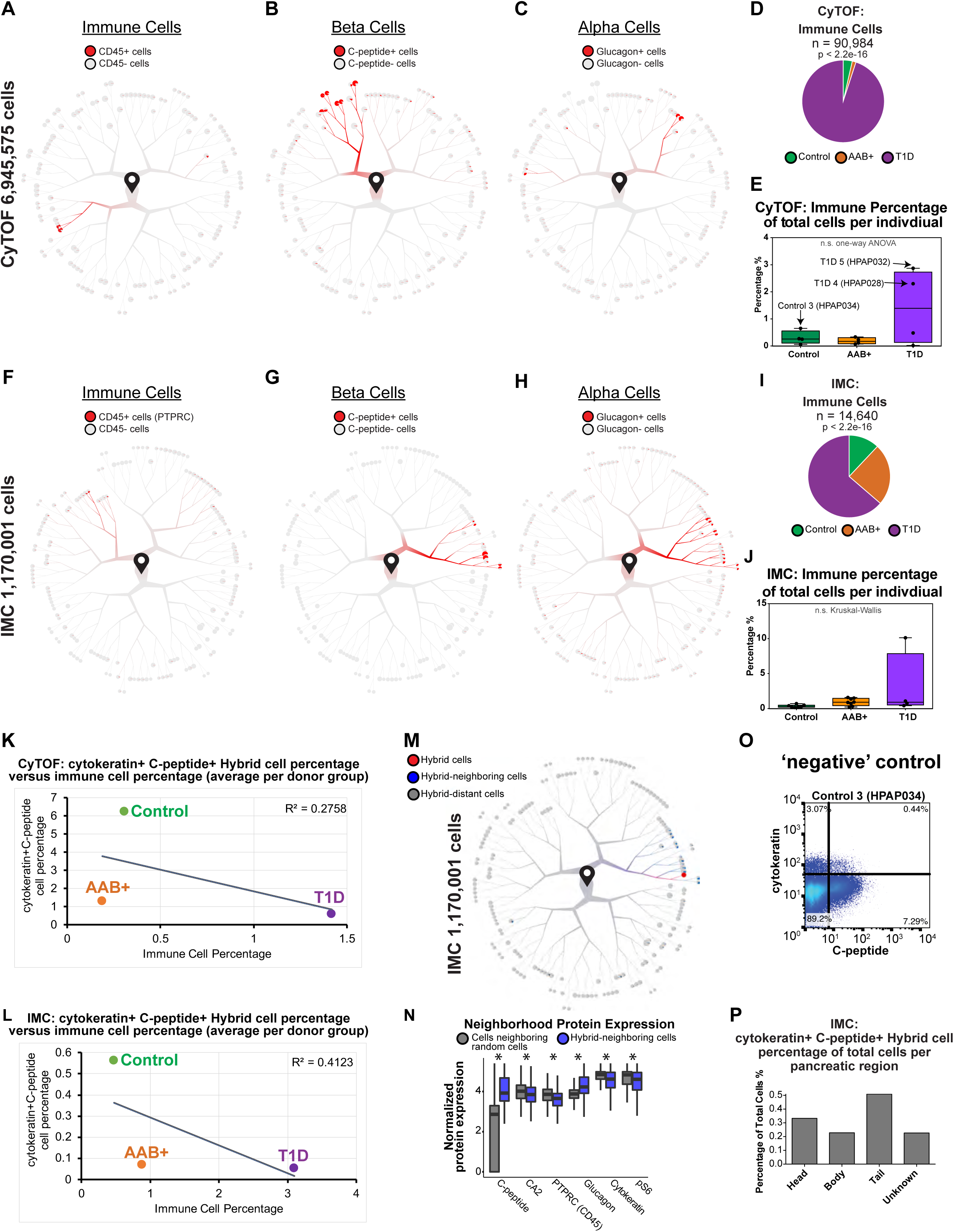
CyTOF and IMC validation of Hybrid cells. (A) Cells begin at the start pin symbol, and from there are partitioned based on similarities and differences in protein expression. Dendrogram visualization of the immune cell cluster, CD45 positive (+) cells, as determined by the analysis of the flow cytometry by time-of-flight (CyTOF) data. (B) Dendrogram visualization of the beta cell cluster, C-peptide positive (+) cells, as determined by the analysis of the CyTOF data. (C) Dendrogram visualization of the alpha cell cluster, Glucagon positive (+) cells, as determined by the analysis of the CyTOF data. (D) Pie chart displaying immune cells and the relative proportions of each donor group from the CyTOF data. P-value presented is the result of the Chi-squared test. (E) Box plots displaying immune cell percentage of total cells per individual across donor groups from the CyTOF data. (F) Dendrogram visualization of the immune cell cluster, CD45 positive (+) cells, as determined by the analysis of the imaging mass cytometry (IMC) data. (G) Dendrogram visualization of the beta cell cluster, C-peptide positive (+) cells, as determined by the analysis of the IMC data. (H) Dendrogram visualization of the alpha cell cluster, Glucagon positive (+) cells, as determined by the analysis of the IMC data. (I) Pie chart displaying immune cells and the relative proportions of each donor group from the IMC data. P-value presented is the result of the Chi-squared test. (J) Box plots displaying immune cell percentage of total cells per individual across donor groups from the IMC data. (K-L) Cytokeratin+ C-peptide+ Hybrid cells versus immune percentage of total cells. For each donor group, the mean immune cell percentage of total cells and the mean Hybrid cell percentage of total cells across all individual donors per donor group was computed to generate one data point per donor group for CyTOF (K) or IMC (L) data. (M) Dendrogram visualization of Hybrid cell (red), Hybrid-Neighboring cell (blue), and Hybrid-Distant cell (grey) enriched clusters as determined by leveraging the spatial architecture provided by IMC data. Dendrograms provided in fig. S17F-H showing immune, beta, and alpha clusters support that Hybrid-neighboring cells are predominately alpha and beta cells, which further supports that Hybrid cells are located within the islets (endocrine compartment) rather than the exocrine compartment of the pancreas. (N) Boxplots showing the normalized protein expression of different canonical markers in Hybrid-Neighboring cells (blue) versus cells neighboring random cells (grey). The number of random cells evaluated was equal to the number of Hybrid cells. Differential marker expression significance for neighbors in the IMC analysis was determined using permutation tests. For each marker, the distribution of that marker value for each of the designated n neighbors was compared against 100 distributions derived from n random cells across the entire IMC tree. * indicates p-value < 0.01. (O) Two parameter CyTOF analysis of cytokeratin and C-peptide expression in single cells from Control donor #3 (HPAP034), a donor with a very low percentage of Hybrid cells as determined by TooManyCells analysis of the CyTOF data. (P) Bar plot displaying the proportion of Hybrid cells from each pancreatic region from the IMC data.

**Figure S18:**
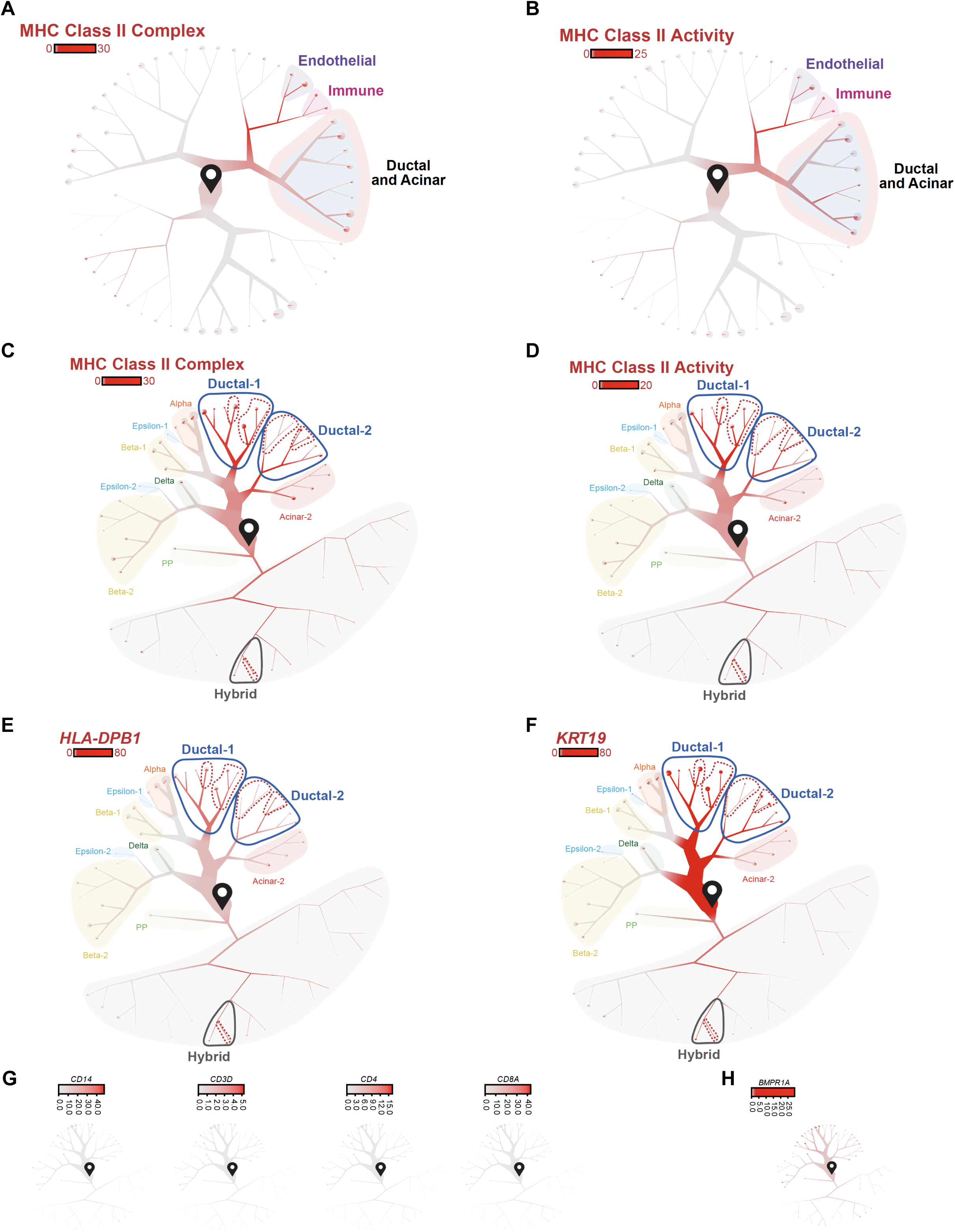
Corroboration of HLA-DR+ Ductal cells. (A-B) Cells begin at the start pin symbol, and from there are partitioned based on similarities and differences in protein expression. Dendrograms highlighting the expression of the MHC class II complex (A) or MHC class II activity (B) across the dendrogram of all cells in Figure 1C. Scale bars represent normalized transcript numbers (mean across all MHC class II complex genes (A) or MHC class II activity genes (B)). (C-D) Dendrograms highlighting the expression of the MHC class II complex (C) or MHC class II activity (D) across the dendrogram of ductal and endocrine cells in Figure 1D. Scale bars represent normalized transcript numbers (mean across all MHC class II complex genes (C) or MHC class II activity genes (D)). (E-F) Dendrograms highlighting the expression of the *HLA-DPB1* (E) or *KRT19* (F) across the dendrogram of ductal and endocrine cells in Figure 1D. Scale bars represent normalized transcript numbers. (G) Dendrograms highlighting the expression of the immune-related genes across the dendrogram of ductal and endocrine cells in Figure 1D. Scale bars represent normalized transcript numbers. (H) Dendrograms highlighting the expression of the *BMPR1A* across the dendrogram of ductal and endocrine cells in Figure 1D. Scale bars represent normalized transcript numbers.

**Figure S19:**
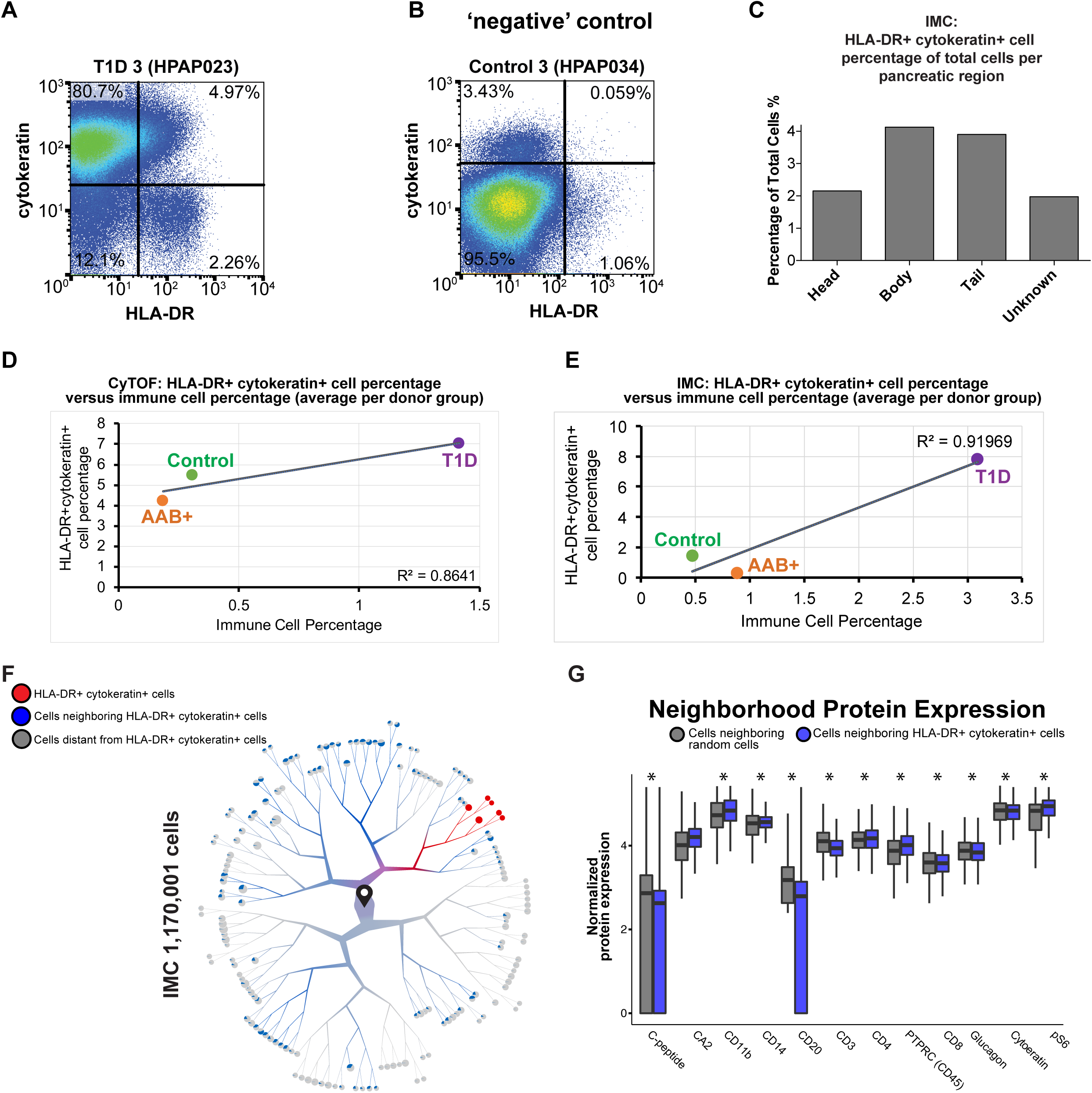
CyTOF and IMC validation of HLA-DR+ Ductal cells. (A) Two parameter CyTOF analysis of HLA-DR and cytokeratin protein expression in single cells from T1D donor #3 (HPAP023). (B) Two parameter CyTOF analysis of HLA-DR and cytokeratin protein expression in single cells from Control donor #3 (HPAP034), a donor with a very low percentage of HLA-DR+ ductal cells as determined by unbiased analysis of CyTOF data with TooManyCells. (C) Bar plot displaying the proportion of HLA-DR+ cytokeratin+ cells from each pancreatic region from the IMC data. (D-E) HLA-DR+ cytokeratin+ cells versus immune percentage of total cells. For each donor group, the mean immune cell percentage of total cells and the mean HLA-DR+ ductal cell percentage of total cells across all individual donors per donor group was computed to generate one data point per donor group for CyTOF (D) or IMC (E) data. (F) Dendrogram visualization of the clusters of HLA-DR+ cytokeratin+ cells (red), cells neighboring HLA-DR+ cytokeratin+ (blue), and cells distant from HLA-DR+ cytokeratin+ cells (grey) as determined by leveraging the spatial architecture provided by IMC data. Cells begin at the start pin symbol, and from there are partitioned based on similarities and differences in protein expression. (G) Boxplots showing the normalized protein expression of different canonical markers in cells neighboring HLA-DR+ cytokeratin+ cells (blue) versus cells neighboring random cells (grey). The number of random cells evaluated was equal to the number of HLA-DR+ cytokeratin+ cells. Differential marker expression significance for neighbors in the IMC analysis was determined using permutation tests. For each marker, the distribution of that marker value for each of the designated n neighbors was compared against 100 distributions derived from n random cells across the entire IMC tree. * indicates p-value < 0.01.

**Figure S20:**
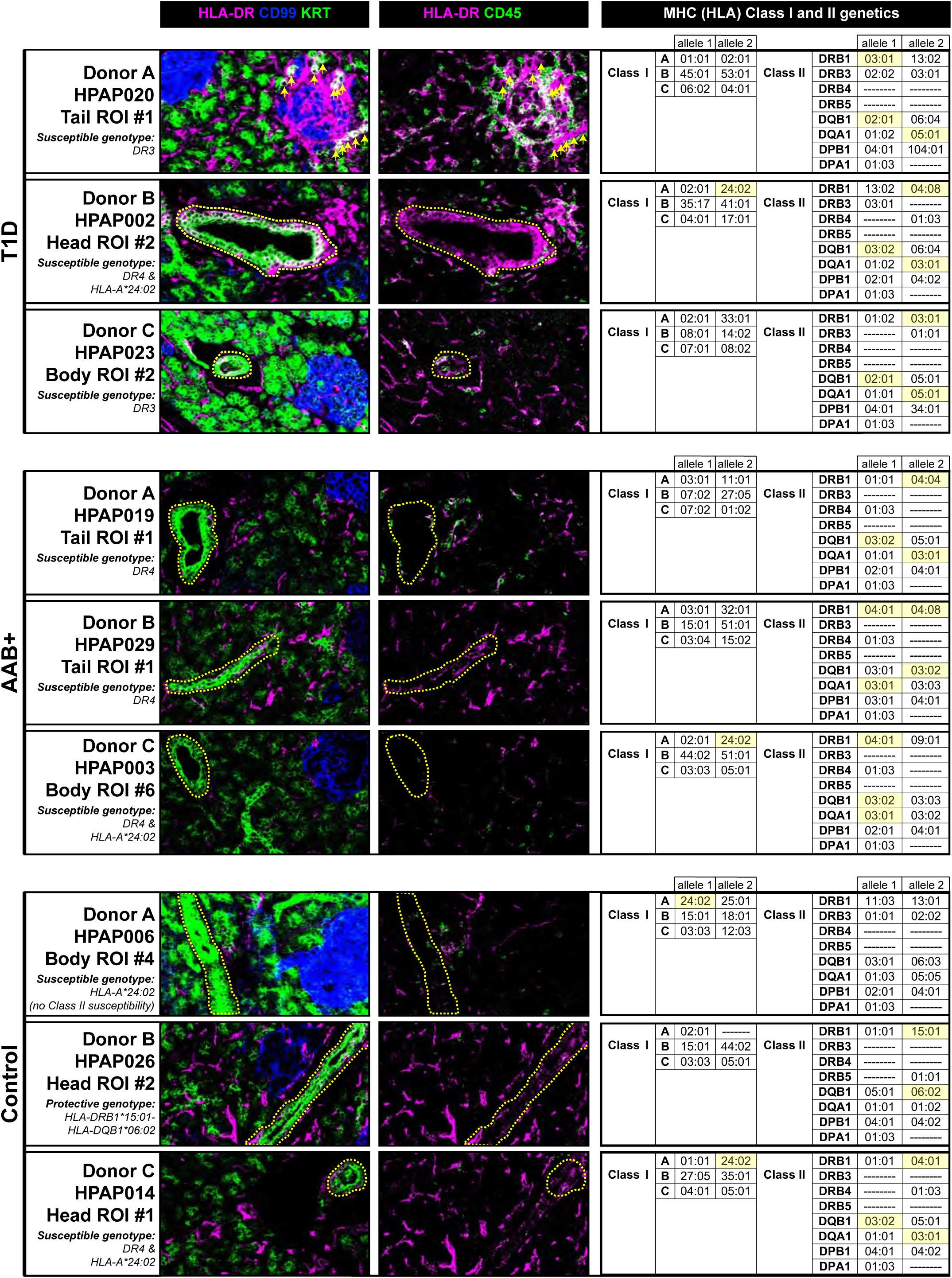
MHC Class II positive ductal cells are present in islets of all donor types, and a high percentage of MHC Class II positive ductal cells in T1D donors appears to be correlated with an inflammatory environment. (Left) Imaging mass cytometry (IMC) in a region of interest (ROI) in pancreatic tissue from three representative individual donors for each donor group type (T1D, AAB+, and Control). HLA-DR is a general marker of MHC Class II (HLA-DR) expression, CD99 is a general islet marker, KRT (pan-keratin) is a ductal cell marker, and CD45 (PTPRC) is a general immune cell marker. Notably, HLA-DR+ ductal cells were primarily located in large ductal structures (outlined in yellow). The images presented here are publicly available at https://www.pancreatlas.org/datasets/508. (Right) HLA typing performed by next generation sequencing. Comprehensive clinical information about each donor is provided in PANC-DB: https://hpap.pmacs.upenn.edu/. Highlighted in yellow are the particular HLA alleles contributing to the susceptible or protective genotypes, which are abbreviated for each donor on the left side of the figure as follows. The four susceptible genotypes assessed were (1) HLA-DRB1*03:01-HLA-DQA1*05:01-HLA-DQB1*02:01 (abbreviated as ‘DR3’, referring to the haplotype bearing the DRB1*03 allele); (2) HLA-DRB1*04:01/02/04/05/08-HLA-DQA1*03:01-HLA-DQB1*03:02/04 (or HLA-DQB1*02) (abbreviated as ‘DR4’, referring to the haplotype bearing the DRB1*04 allele); (3) HLA-A*24:02; and (4) HLA-B*39:06. The two protective genotypes assessed were (1) HLA-DRB1*15:01-HLA-DQB1*06:02 and (2) HLA-DRB1*07:01-HLA-DQB1*03:03 (Howson et al., 2012; Howson et al., 2009; Inshaw et al., 2020; Nejentsev et al., 2007; Noble and Valdes, 2011; Valdes et al., 2012). Notably, HLA-DR+ ductal cells were found across all HLA genotypes, including both susceptible and protective genotypes.

## Tables

**Table 1:** Donor clinical information

**Table 2:** Differential gene list between Beta-1 cells (positive fold change (FC)) and Beta-2 cells (negative FC)

**Table 3:** Differential gene list between Epsilon-1 cells (positive fold change (FC)) and Epsilon-2 cells (negative FC)

**Table 4:** Differential gene list between Ductal-1 cells (positive fold change (FC)) and Ductal-2 cells (negative FC)

**Table 5:** Differential gene list between Ductal-2 cells (positive fold change (FC)) and Acinar-2 cells (negative FC)

**Table 6:** Differential gene list between AAB+ Beta-1 cells (positive fold change (FC)) and Control Beta-1 cells (negative FC)

**Table 7:** Differential gene list between T1D Beta-1 cells (positive fold change (FC)) and Control Beta-1 cells (negative FC)

**Table 8:** Differential gene list between AAB+ Beta-2 cells (positive fold change (FC)) and Control Beta-2 cells (negative FC)

**Table 9:** Differential gene list between T1D Beta-2 cells (positive fold change (FC)) and Control Beta-2 cells (negative FC)

**Table 10:** Differential gene list between AAB+ Ductal-1 cells (positive fold change (FC)) and Control Ductal-1 cells (negative FC)

**Table 11:** Differential gene list between T1D Ductal-1 cells (positive fold change (FC)) and Control Ductal-1 cells (negative FC)

**Table 12:** Differential gene list between AAB+ Ductal-2 cells (positive fold change (FC)) and Control Ductal-1 cells (negative FC)

**Table 13:** Differential gene list between T1D Ductal-2 cells (positive fold change (FC)) and Control Ductal-1 cells (negative FC)

**Table 14:** Differential gene list between AAB+ Acinar-1 cells (positive fold change (FC)) and Control Acinar-1 cells (negative FC)

**Table 15:** Differential gene list between T1D Acinar-1 cells (positive fold change (FC)) and Control Acinar-1 cells (negative FC)

**Table 16:** Differential gene list between AAB+ Acinar-2 cells (positive fold change (FC)) and Control Acinar-2 cells (negative FC)

**Table 17:** Differential gene list between T1D Acinar-2 cells (positive fold change (FC)) and Control Acinar-2 cells (negative FC)

**Table 18:** Differential gene list between AAB+ Alpha cells (positive fold change (FC)) and Control Alpha cells (negative FC)

**Table 19:** Differential gene list between T1D Alpha cells (positive fold change (FC)) and Control Alpha cells (negative FC)

**Table 20:** Differential gene list between AAB+ Delta cells (positive fold change (FC)) and Control Delta cells (negative FC)

**Table 21:** Differential gene list between T1D Delta cells (positive fold change (FC)) and Control Delta cells (negative FC)

**Table 22:** Differential gene list between AAB+ PP cells (positive fold change (FC)) and Control PP cells (negative FC)

**Table 23:** Differential gene list between T1D PP cells (positive fold change (FC)) and Control PP cells (negative FC)

**Table 24:** Differential gene list between AAB+ Endothelial cells (positive fold change (FC)) and Control Endothelial cells (negative FC)

**Table 25:** Differential gene list between T1D Endothelial cells (positive fold change (FC)) and Control Endothelial cells (negative FC)

**Table 26:** Differential gene list between AAB+ Stellate cells (positive fold change (FC)) and Control Stellate cells (negative FC)

**Table 27:** Differential gene list between T1D Stellate cells (positive fold change (FC)) and Control Stellate cells (negative FC)

**Table 28:** Differential gene list between AAB+ Hybrid cells (positive fold change (FC)) and Control Hybrid cells (negative FC)

**Table 29:** Differential gene list between T1D Hybrid cells (positive fold change (FC)) and Control Hybrid cells (negative FC)

**Table 30:** Differential gene list between AAB+ Epsilon-1 cells (positive fold change (FC)) and Control Epsilon-1 cells (negative FC)

**Table 31:** Differential gene list between T1D Epsilon-1 cells (positive fold change (FC)) and Control Epsilon-1 cells (negative FC)

**Table 32:** Differential gene list between AAB+ Epsilon-2 cells (positive fold change (FC)) and Control Epsilon-2 cells (negative FC)

**Table 33:** Differential gene list between T1D Epsilon-2 cells (positive fold change (FC)) and Control Epsilon-2 cells (negative FC)

**Table 34:** Differential gene list between AAB+ Immune cells (positive fold change (FC)) and Control Immune cells (negative FC)

**Table 35:** Differential gene list between T1D Immune cells (positive fold change (FC)) and Control Immune cells (negative FC)

**Table 36:** Differential gene list between T1D Beta-1 cells (positive fold change (FC)) and AAB+ Beta-1 cells (negative FC)

**Table 37:** Differential gene list between T1D Beta-2 cells (positive fold change (FC)) and AAB+ Beta-2 cells (negative FC)

**Table 38:** Differential gene list between T1D Ductal-1 cells (positive fold change (FC)) and AAB+ Ductal-1 cells (negative FC)

**Table 39:** Differential gene list between T1D Ductal-2 cells (positive fold change (FC)) and AAB+ Ductal-2 cells (negative FC)

**Table 40:** Differential gene list between T1D Acinar-1 cells (positive fold change (FC)) and AAB+ Acinar-1 cells (negative FC)

**Table 41:** Differential gene list between T1D Acinar-2 cells (positive fold change (FC)) and AAB+ Acinar-2 cells (negative FC)

**Table 42:** Differential gene list between T1D Alpha cells (positive fold change (FC)) and AAB+ Alpha cells (negative FC)

**Table 43:** Differential gene list between T1D Delta cells (positive fold change (FC)) and AAB+ Delta cells (negative FC)

**Table 44:** Differential gene list between T1D PP cells (positive fold change (FC)) and AAB+ PP cells (negative FC)

**Table 45:** Differential gene list between T1D Endothelial cells (positive fold change (FC)) and AAB+ Endothelial cells (negative FC)

**Table 46:** Differential gene list between T1D Stellate cells (positive fold change (FC)) and AAB+ Stellate cells (negative FC)

**Table 47:** Differential gene list between T1D Hybrid cells (positive fold change (FC)) and AAB+ Hybrid cells (negative FC)

**Table 48:** Differential gene list between T1D Epsilon-1 cells (positive fold change (FC)) and AAB+ Epsilon-1 cells (negative FC)

**Table 49:** Differential gene list between T1D Epsilon-2 cells (positive fold change (FC)) and AAB+ Epsilon-2 cells (negative FC)

**Table 50:** Differential gene list between T1D Immune cells (positive fold change (FC)) and AAB+ Immune cells (negative FC)

**Table 51:** Differential gene list between Endocrine-like Hybrid cells (positive fold change (FC)) and beta cells (both Beta-1 and Beta-2 cells; negative FC)

**Table 52:** Differential gene list between Endocrine-like Hybrid cells (positive fold change (FC)) and Alpha cells (negative FC)

**Table 53:** Differential gene list between Ductal-like Hybrid cells (positive fold change (FC)) and beta cells (both Beta-1 and Beta-2 cells; negative FC)

**Table 54:** Differential gene list between Ductal-like Hybrid cells (positive fold change (FC)) and ductal cells (both Ductal-1 and Ductal-2 cells; negative FC)

**Table 55:** Differential gene list between Acinar-like Hybrid cells (positive fold change (FC)) and beta cells (both Beta-1 and Beta-2 cells; negative FC)

**Table 56:** Differential gene list between Acinar-like Hybrid cells (positive fold change (FC)) and Acinar-2 cells (negative FC)

**Table 57:** Differential gene list between all Hybrid cells (positive fold change (FC)) and all remaining cells in the ductal and endocrine tree (negative FC)

**Table 58:** CyTOF Panel

**Table 59:** IMC Panel

**Table 60:** Garnett cell type marker file (*37*)

